# VlbZIP30 of grapevine functions in drought tolerance via the abscisic acid core signaling pathway

**DOI:** 10.1101/231811

**Authors:** Mingxing Tu, Xianhang Wang, Yanxun Zhu, Dejun Wang, Xuechuan Zhang, Ye Cui, Yajuan Li, Min Gao, Zhi Li, Xiping Wang

## Abstract

Drought stress limits the growth and development of grapevines, thereby reducing productivity, but the mechanisms by which grapevines respond to drought stress remain largely uncharacterized. Here, we characterized a group A bZIP gene from ‘Kyoho’ grapevine, *VlbZIP30,* which was shown to be induced by abscisic acid (ABA) and dehydration stress. Overexpression of *VlbZIP30* in transgenic Arabidopsis enhanced dehydration tolerance during seed germination, and in the seedling and adult stages. Transcriptome analysis revealed that a major proportion of ABA- and/or drought-responsive genes are transcriptionally regulated by *VlbZIP30* during ABA or mannitol treatment at the cotyledon greening stage. We identified an *A. thaliana* G-box motif (CACGTG) and a potential grapevine G-box motif (MCACGTGK) in the promoters of the 39 selected *A. thaliana* genes up-regulated in the transgenic plants and in the 35 grapevine homologs, respectively. Subsequently, using two grapevine-related databases, we found that 74% and 84% (a total of 27 genes) of the detected grapevine genes were significantly up-regulated by ABA and drought stress, respectively, suggesting that these 27 genes involve in ABA or dehydration stress and may be regulated by *VlbZIP30* in grapevine. We propose that *VlbZIP30* functions as a positive regulator of drought-responsive signaling in the ABA core signaling pathway.

**Highlight:** *VlbZIP30* positively regulate plant drought tolerance through regulated the expression of 27 grapevine candidate genes via G-box *cis*-element (MCACGTGK) in ABA signaling pathway.

## Introduction

Grapevines are amongst the world’s major fruit crops, and their fruits can be consumed either fresh or dried or be processed into wines, spirits and vinegar, or transformed into pharmaceutical products that promote human health (Pilati *et al.*, 2017). However, abiotic stress, such as drought, perturb the metabolism and growth of grapevines, leading to a loss of yield and reduced fruit quality (Ferreira *et al.*, 2004). Consequently, increasing the resistance of grapevines to drought stress is an important factor in ensuring yield stability.

Stress signaling in plants can be transduced by various signaling components, including second messengers (e.g. Ca^2+^), signal transduction factors, including protein kinases and phosphatases, hormonessuch as abscisic acid (ABA), and transcription factors (TFs). Such signaling associated with drought has been shown to cause changes in physiological, morphological and molecular processes, including the activation of many drought stress-related genes and the accumulation of a range of proteins, reflecting a drought stress response (Zhu, 2002; Yamaguchi-Shinozaki and Shinozaki, 2006; Lata and Prasad, 2011; Tang *et al.*, 2012).

ABA is considered to be a stress hormone and has been particularly associated with drought tolerance, although it is also involved in various developmental processes, including seed germination, seedling growth and development (Finkelstein *et al.*, 2002; Tang *et al.*, 2012). In the context of a drought response it has been shown to mediate stomatal closure and to promote cuticular wax biosynthesis (Nambara and Kuchitsu, 2011). Analyses of the underlying molecular mechanisms have demonstrated that both ABA-dependent and ABA-independent pathways are involved in drought stress responses (Shinozaki and Yamaguchi-Shinozaki, 2000; Yamaguchi-Shinozaki and Shinozaki, 2006). ABA-mediated drought tolerance involves complex signaling networks, the core components of which have been identified. Briefly, when ABA is present, it binds to the ABA receptors PYR/PYL/RCAR (PYRABACTIN RESISTANCE1/PYR1 -like/REGULATORY COMPONENT OF ABA RECEPTOR1), which interact with the PP2C (PROTEIN PHOSPHATASE 2C) proteins, forming a complex and releasing the inhibitory effect of PP2Cs on SnRK2 (SUCROSE-NONFERMENTING1-RELATED PROTEIN KINASE2) protein kinases. The activated SnRK2s proteins subsequently phosphorylate different downstream TFs, such as AREB1 (ABA-RESPONSE-ELEMENT BINDING1) and ABI5 (ABA INSENSITIVE5), which regulate the expression of ABA-responsive genes (Fujii and Zhu, 2009; Fujita *et al.*, 2009; Ma *et al.*, 2009; Nakashima *et al.*, 2009; Park *et al.*, 2009; Danquah *et al.*, 2014).

TFs are generally identified according to conserved sequences, known as the DNA-binding domains. One of the largest TF families in higher plants is the bZIP family, members of which are characterized by a basic region/leucine zipper domain (Van Leene *et al.*, 2016). Previous studies have shown that bZIP proteins function as regulators of signaling networks by specifically binding *cis*-elements containing an core ACGT, such as the ABA-responsive element (ABRE; PyACGTGGC), the G-box (CACGTG) and the C-box (GACGTC) (Yamaguchi-Shinozaki *et al.*, 1990; Foster *et al.*, 1994), in the promoters of their target genes, to either activate or repress their expression (Mitsuda and Ohme-Takagi, 2009).

A number of studies have shown that bZIP TFs are important regulators of drought stress signaling in the ABA-dependent pathway, mostly in association with seed germination and post-germination growth. The involvement of bZIP TFs *(ABF1, AREB1/ABF2, ABF3, AREB2/ABF4)* in the regulation of drought responses was first reported in the model plant *Arabidopsis thaliana* (Kang *et al.*, 2002; Kim *et al.*, 2004; Fujita *et al.*, 2005; Yoshida *et al.*, 2010; Yoshida *et al.*, 2015). Following these studies, drought-related bZIP genes have been identified in a range of other species, including *OsABI5* in rice (*Oryza sativa*) (Zou *et al.*, 2008), *LIP19* in wheat (*Triticumaestivum*) (Kobayashi *et al.*, 2008), *ABP9* in maize (Zea *mays*) (Zhang *et al.*, 2011), *SlAREB1* in tomato (*Solanum lycopersicum*) (Orellana *et al.*, 2010), *GmbZIP1* in soybean (*Glycine max*) (Gao *et al.*, 2011), *ThbZIP1* in *Tamarixhispida* (Ji *et al.*, 2013), *CaBZ1* in hot pepper (*Capsicum annuum*) (Moon *et al.*, 2015), and *PtrABF* in trifoliate orange (*Citrus trifoliata*) (Zhang *et al.*, 2015).

In order to improve the typically poor drought resistance of grapevines, researchers have focused their attention on the identification of drought-related TFs. Several, such as *CBF1/2/3/4* (Xiao *et al.*, 2006; Siddiqua and Nassuth, 2011; Li *et al.*, 2013), *WRKY11* (Liu *et al.*, 2011a), *ERF1/2/3* (Zhu *et al.*, 2013), *NAC26* (Fang *et al.*, 2016), and *PAT1* (Yuan *et al.*, 2016), have been identified and their overexpression in *A. thaliana* has been shown to enhance drought resistance. However, to date, only a few grapevine bZIP TFs have been functionally characterized during a drought stress response (Gao *et al.*, 2014; Tu *et al.*, 2016a, b), and their regulatory mechanisms are not well understood.

In this current study, we cloned a group A bZIP TF, *VlbZIP30*, from ‘Kyoho’ grapevine (*Vitis labrusca*χ *V. vinifera*) and ectopically expressed it in *A. thaliana.* The results of physiological and transcriptomic analyses of the transgenic lines are presented and its putative function in drought-responsive signaling via the ABA signaling pathway in grapevine is discussed.

## Materials and methods

### Plant material and growth conditions

The two-year-old ‘Kyoho’ grapevine (*Vitis labrusca×V vinifera*) plants used in this study were grown in the grapevine repository of the Northwest A&F University, Yangling, Shaanxi, China. *A. thaliana* ecotype Columbia (Col-0) plants used as both wild type (WT) and for transgenic experiments were grown in a greenhouse at 21°C under long-day (LD) conditions (16 h light/8 h dark).

### Dehydration stress and ABA treatment of grapevine leaves

For dehydration treatments, grapevine shoots with three well-developed leaves were detached and immediately placed on dry filter paper in an illumination incubator at 25°C, with a relative humidity of 60-70%, under LD conditions (16 h light/8 h dark). For ABA treatments, leaves were sprayed with 100 μM ABA while the shoots were immersed in water, and the plants were then placed under the same ambient conditions as above. Leaves from the same position were collected from three independent replicates of each treatment at 1, 2, 4, 6, 9, 12, and 24 h after initiating treatment. The 0 h samples were collected before each treatment was initiated and used as control samples. All samples were immediately frozen in liquid nitrogen and stored at –80°Cuntil further analysis.

### Bioinformatic analysis

Full-length amino acid sequences of bZIP TFs from *A. thaliana* and grapevine were obtained from The Arabidopsis Information Resource (TAIR; http://www.arabidopsis.org/index.jsp) and EnsemblPlants (http://plants.ensembl.org/index.html), respectively. Multiple amino acid sequence alignments were generated using DNAMAN software (Version 5.2.2.0, LynnonBiosoft, USA) with default parameters, and a phylogenetic tree was constructed using the neighbor-joining (NJ) method and MEGA software (version 5.05), with 1,000 bootstrap replicates, as previously described (Tu *et al.*, 2016b). The predicted phosphorylation sites (C1, C2, C3, and C4) and highly conserved bZIP domain were analyzed as previously described (Fujita *et al.*, 2005).

### Transformation and characterization of transgenic plants

The plant transformation vectors 35S: *VlbZIP30* and Pro_*VlbZIP30*_-*GUS* (β-glucosidase, details of vector construction are supplied in Supplementary Method S1) were transformed into *A. thaliana* by the floral dip method using *Agrobacterium tumefaciens* (strain GV3101) (Clough and Bent, 1998).

For each construct, seeds of the T0 and T1 plants were screened on Murashige-Skoog (MS) agar medium supplemented with 100 mg/L kanamycin. For phenotypic investigation, the three T3 homozygous lines (OE1, OE6 and OE23) with the highest levels of *VlbZIP30* expression, were used. To assess the expression of *GUS* in the Pro*_VlbZIP30_:GUS* transgenic plants, T3 homozygous lines from 3 independent transgenic lines were analyzed. Seeds from each of the three selected T3 homozygous lines and from WT plants were vernalized and sterilized as previously described (Tu *et al.*, 2016b).

### Histochemical GUS assay

An *in situ* GUS activity assay was performed as previously described (Tu *et al.*, 2016b).

### Osmotic stress and ABA treatment of transgenic seedlings

WT and transgenic seeds were harvested at the same time. For seed germination and cotyledon greening analyses, approximately 100 seeds from WT and each 35S:*VlbZIP30* line (OE1, OE6 and OE23) were grown on MS agar medium, MS agar medium containing 300mM or 350 mM mannitol, or on MS agar medium containing 0.5 μM or 1 μM ABA, at 21°C with a 16 h light/8 h dark cycle. Germination and cotyledon greening rates were defined as the obvious emergence of the seedling radicle through the seed coat and green coloration of cotyledons, respectively (Tu *et al.*, 2016b). The seedlings were sampled after counting to measure the endogenous ABA contents.

For the osmotic stress and ABA treatments, 7-day-old WT and transgenic seedlings were transferred from MS medium plates into MS agar medium, or MS agar medium supplemented with 300 mM or 350 mM mannitol, or MS agar medium supplemented with 50 μM or 100 μM ABA. The root lengths were measured 7 d after the transfer.

### Transcriptome analysis and identification of differentially expressed genes (DEGs)

Seeds from WT and transgenic lines were cultivated on MS agar medium, with or without stress treatment (0.5 μM ABA or 300 mM mannitol) for 7 d, and collected for RNA extraction. For each RNA purification biological replicate, 300 seedlings of WT or the OE lines from three MS agar plates were pooled to form a single sample. Three independent RNA samples were used for each experiment.

Total RNA was extracted using the E.Z.N.A. Plant RNA Kit (Omega Bio-tek, USA, R6827-01) according to the manufacturer’s protocol (Invitrogen). RNA concentration and integrity were confirmed using a NanoDrop 2000 spectrophotometer (Thermo Fisher Scientific, Wilmington, DE, USA) and an Agilent 2100 Bioanalyzer (Agilent Technologies, CA, USA). The construction of RNA-Seq libraries and sequencing were performed by the Biomarker Biotechnology Corporation (Beijing, China). The libraries were generated using the NEBNext UltraTM RNA Library Prep Kit for Illumina (NEB, USA) following the manufacturer’s recommendations. Sequencing of the purified libraries was carried out using an Illumina HiseqXten platform (Illumina, NEB, USA) generating paired-end reads. The raw reads were cleaned by removing reads containing adapter sequences, reads containing poly-N and low quality reads. The cleaned reads from each sample were aligned to the *A. thaliana* reference genome from TAIR using the Tophat2 software (Kim *et al.*, 2013). Gene expression levels were determined by fragments per kilobase of transcript per million fragments mapped (FPKM), and the DEGs were identified using edgeR software (Robinson *et al.*, 2010), with a threshold of false discovery rate (FDR)<0.05 and absolute log2FC (fold change)>1. All raw sequence data in this study have been submitted to the NCBI Short Read Archive (SRA) under BioProject accession number PRJNA419694.

### Transcriptome data analysis

The Venn diagrams were made using the BMK Cloud platform (www.biocloud.net). Annotations for DEGs were retrieved from TAIR. Gene Ontology (GO) enrichment analyses were performed for the functional categorization of DEGs based on the PageMan profiling tool (Usadel *et al.*, 2006) and Arabidopsis Functional Modules Supporting Data (Heyndrickx and Vandepoele, 2012). The grapevine orthologs of the *A. thaliana* genes were identified using TBLASTX software (Altschul *et al.*, 1997) with the highest score. Motif predictions were performed using the promoter region 1,500bp upstream of the start codons of the *A. thaliana* (AT) and grapevine (VIT) genes using DREME software (http://meme-suite.org/tools/dreme). The heat maps were constructed using HemI software (Deng *et al.*, 2014). To identify the predicted grapevine genes, two grapevine-related ABA (Pilati *et al.*, 2017) and drought stress (Rocheta *et al.*, 2016) databases were downloaded from the National Center for Biotechnology Information (NCBI) under BioProject accession number PRJNA369777 and the Gene Expression Omnibus (GEO) database under the number GSE57669.

### Water loss assay and drought treatment of mature transgenic seedlings

For the water loss assay, rosette leaves of 3-week-old WT and transgenic plants were detached and immediately placed on dry filter paper. The samples, together with the paper, were then placed in the laboratory at ambient temperature with a relative humidity of 45-50%, and weighed at the indicated times. The fresh weight of the leaves was measured every 30 min to calculate relative water loss. The leaves were sampled after dehydration to examine cell death phenotypes, measure endogenous ABA content, antioxidant enzyme activity, and levels of ROS. The 0 h samples collected before dehydration were used as the negative control. For the drought treatment, plants were initially grown for 3 weeks under a normal watering regime and then water was withheld for 8 d. Survival rates were scored after re-watering for 3 days. Well-watered plants were used as the negative control.

### Analysis of electrolyte leakage, MDA content, cell death, reactive oxygen species (ROS) levels, antioxidant enzyme activity and ABA content

The relative electrolyte leakage, MDA content and antioxidant enzyme activity were measured as previously described (Tu *et al.*, 2016b), as was ABA content (Tu *et al.*, 2016a). Histochemical staining procedures were used to detect *in situ* reactive oxygen species (ROS) levels and dead cells as previously described (Tu *et al.*, 2016b).

### Stoma tal aperture analysis

Stomatal aperture assays were performed as previously described (Tu *et al.*, 2016b).

### RNA extraction and quantitative real-time PCR (qRT-PCR)

Total RNA was extracted from the grapevine leaves at 8 time points (0, 1, 2, 4, 6, 9, 12, and 24 h) after ABA and dehydration treatment using the E.Z.N.A._Plant RNA Kit (Omega Bio-tek, USA, R6827-01) following the manufacturer’s instructions, as was total *A. thaliana* RNA from the leaves of 3-week-old WT and OE lines collected before and after dehydration. The qRT-PCR analyses were conducted using SYBR Premix Ex Taq II (TliRNaseH Plus) (TaKaRa Biotechnology) and a StepOnePlus™ RT-PCR instrument from Thermo Fisher Scientific with the following thermal profile: 95°C for 30 s, 45 cycles of 95°C for 5 s, and 60°C for 30 s. The expression levels of the grape *ACTIN1* (VIT_04s0044g00580) or *A. thaliana ACTIN2* (AT3G18780) genes were used as references. The specific primers for qRT-PCR are listed in Supplementary Table S1. Relative expression levels were analyzed using the StepOnePlus software (v. 2.3) and the Normalized Expression Method.

## Statistical analysis

Data analysis was performed using Microsoft Excel (Microsoft Corporation, USA). The data were plotted using Sigmaplot (v. 10.0, Systat Inc., CA, USA). Paired t tests were performed to assess significant differences using the SPSS Statistics 17.0 software (IBM China Company Ltd., Beijing, China). All experiments were repeated three times as independent analyses.

## Results

### Identification of VlbZIP30, a group A bZIP TF from grapevine

The *VlbZIP30* (VIT_13s0175g00120) cDNA is 978 bp long and encodes a protein of 325 amino acids. Amino acid sequence analysis showed that, in common with the 8 members of the *A. thaliana* ABF/DPBF bZIP subfamily, VlbZIP30 also contains a leucine zipper domain (Jakoby *et al.*, 2002) and conserved domains predicted as phosphorylation sites (C1, C2, C3 and C4) involved in drought stress or ABA signaling (Fujita *et al.*, 2005) (Fig. 1). A phylogenetic analysis indicated that VlbZIP30 is most closely related to the group A ABF/DPBF TFs, which have previously been shown to be involved in ABA and drought stress signaling in *A. thaliana* (Kang *et al*, 2002; Kim *et al.*, 2002; Fujita *et al.*, 2005; Yoshida *et al*, 2015), and in grapevine (Nicolas *et al.*, 2014; Tu *et al.*, 2016a) (Fig. 2A).

**Fig. 1.**
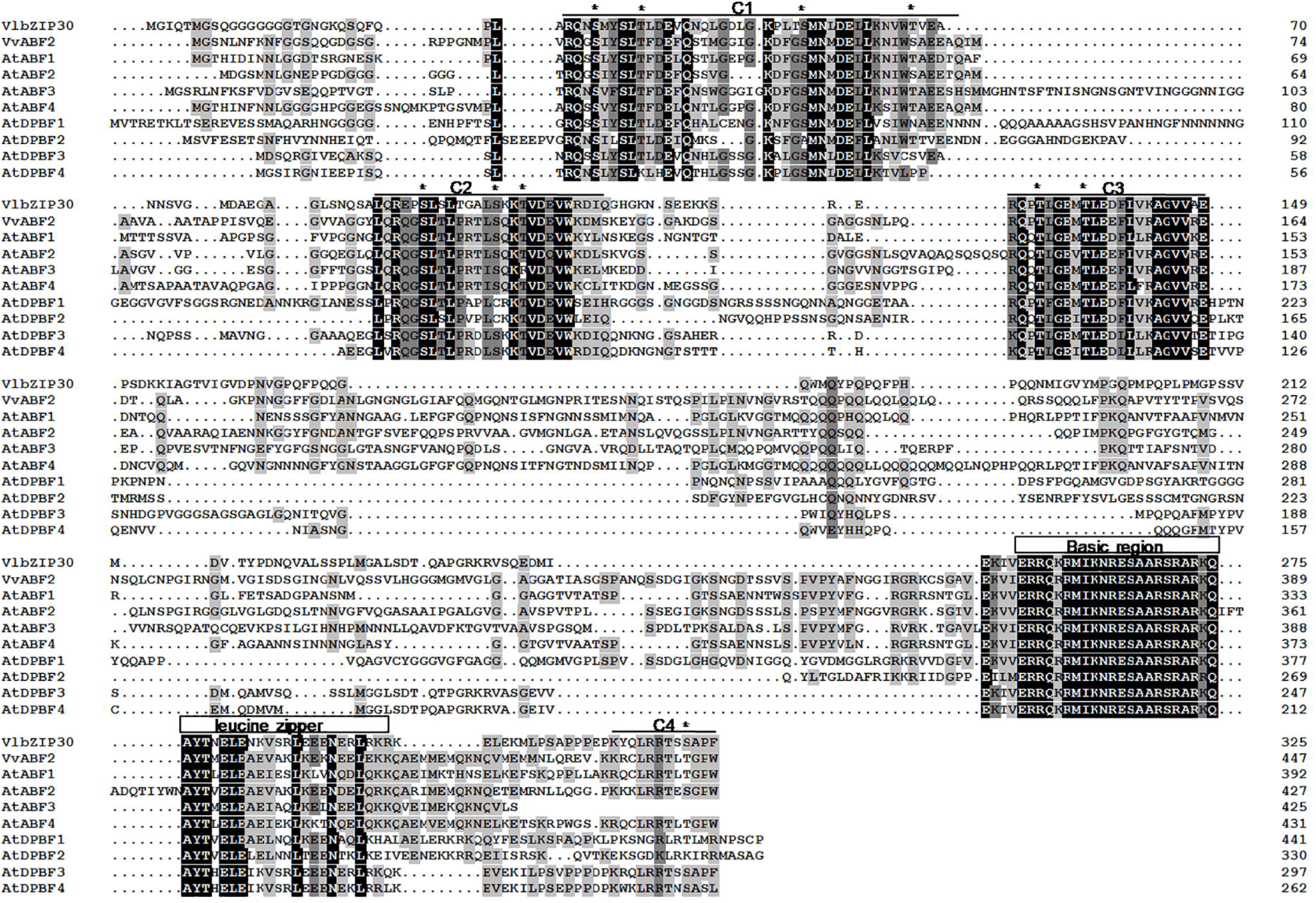
Multiple sequence alignment. Full-length sequence comparison of VlbZIP30 (VIT_13s0175g00120) with the ABF/DPBF subfamily of group A bZIP proteins from *Arabidopsis thaliana* (AT) and grapevine (*Vitis vinifera*, VIT). AtABF1 (AT1G49720), AtABF2 (AT1G45249), AtABF3 (AT4G34000), AtABF4 (AT3G19290), AtDPBF1 (AT2G36270), AtDPBF2 (AT3G44460), AtDPBF3 (AT3G56850), AtDPBF4 (AT2G41070) and VvABF2 (VIT_18s0001g10450) were used for the alignment. Conserved residues with 100%, 75-99 % or 33-75% amino identity are shaded in black, dark gray and light gray, respectively. The conserved bZIP domains are indicated with black rectangles. Putative phosphorylation sites (C1, C2, C3 and C4, underlined) are marked with asterisks.

**Fig. 2.**
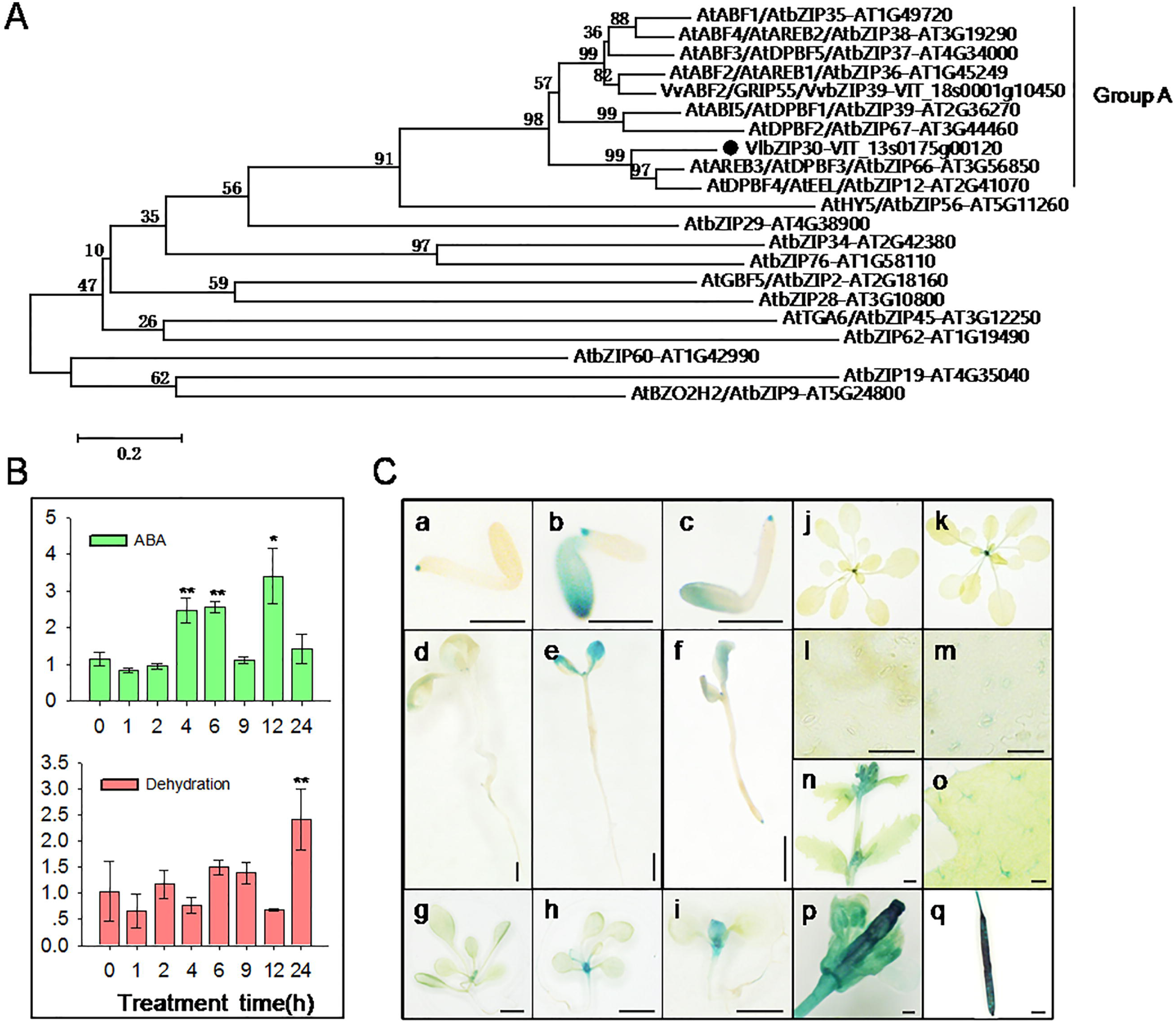
Phylogenetic analysis of VlbZIP30 and expression analysis of *VlbZIP30*. (A) The phylogenetic tree represents VlbZIP30 (black circle) and other bZIP amino acid sequences from *Arabidopsis thaliana* (AT) and grapevine (*Vitis vinifera*, VIT). The clustering of the group A bZIP proteins and the other groups of bZIP proteins (group H, AtHY5; I, AtbZIP29; E, AtbZIP34; L, AtbZIP76; S AtGBF5; B, bZIP28; D AtTGA6; J, AtbZIP62; K, AtbZIP60; F, AtbZIP19; and C, AtBZO2H2) have previously beenreported by Jakoby *et al.* (2002) and Nicolas *et al.* (2014). (B) Expression profiles of *VlbZIP30* in grapevine following abscisic acid (ABA) and dehydration treatments. Data represent the mean values ±SE from three independent experiments. Asterisks indicate statistical significance (*0.01 < P < 0.05, **P < 0.01, Student’s *t*-test) between the treated and untreated control plants. (C) Patterns of *VlbZIP30* promoter-driven GUS (β-glucosidase) expression in *A. thaliana* at different growth stages. Mature embryos cultivated on Murashige-Skoog (MS) agar medium (a), or MS agar medium supplemented with 300 mM mannitol (b) or 0.5 μM ABA (c) for 2 d. Scale bar = 500 μm. Seven-day-old seedlings cultivated on MS agar medium (d), or MS agar medium supplemented with 300 mM mannitol (e) or 0.5 μM ABA (f) for 7 d. Scale bar = 500 μm. Fourteen-day-old seedlings transferred from MS medium plates into MS agar medium (g) or MS agar medium supplemented with 300 mM mannitol (h) or 100 μM ABA (i) for 7 d. Scale bar = 2 mm. (j) 3-week-old plant. (k) 3-week-old plant after dehydration for 2 h. (l) Guard cells of 3-week-old plant. Scale bar = 50 μm. (m) Guard cells of 3-week-old plant after dehydration for 2 h. Scale bar = 50 μm. (n) Inflorescence. Scale bar = 2 mm. (o) Leaf. Scale bar = 200 μm. (p) Flower. Scale bar = 200 μm. (q) Silique. Scale bar = 2 mm.

### Expression of VlbZIP30 is induced by drought and ABA treatment

To test whether *VlbZIP30* is involved in ABA and drought stress signaling, we first evaluated the expression levels of *VlbZIP30* in grapevine following ABA or dehydration treatments using qRT-PCR. As shown in Fig. 2B, ABA caused an increase in *VlbZIP30* expression at 4 h and 6 h shortly after initiation of the treatment. The expression peaked at 12 h, before decreasing for the next 24 h. Dehydration caused an increase in *VlbZIP30* expression at 24 h.

Next, to investigate the temporal and spatial expression patterns of *VlbZIP30* in more detail, histochemical GUS reporter experiments were performed with plants grown under ABA and dehydration stress, as well as under normal conditions. Low levels of GUS staining were observed in 2-d seeds at the germination stage, as well as in 7 d and 14 d old seedlings (Fig. 2C, a, d and g), and GUS activity was significantly enhanced after mannitol (Fig. 2C, b, e and h) and ABA (Fig. 2C, c, f and i) treatments at the same stages. In mature plants, GUS staining was obviously detected in stems, trichomes, flowers and siliques (Fig. 2C, n-q), while only slight staining was detected in leaf petioles (Fig. 2C, j), and no staining was detected in guard cells (Fig. 2C, l). However, after dehydration for 2 h, the leaf petioles and guard cells showed an increase in GUS staining in 3-week-old plants (Fig. 2C, k, m). The dehydration treatments had no effect on the size of the stomatal aperture. These results suggest that *VlbZIP30* expression is regulated by ABA and drought stress.

### Overexpressing VlbZIP30 in A. thaliana reduces mannitol and ABA sensitivity during seed germination and post-germination growth

Three homozygous transformed lines (OE1, OE6 and OE23) with the highest levels of *VlbZIP30* expression were selected based on qRT-PCR analysis (Supplementary Fig. S1). Sterilized seeds of the transgenic lines and WT plants were cultivated on MS agar medium, or MS agar medium containing 300 mM or 350 mM mannitol, or 0.5 μM or 1 μM ABA. The seed germination rates of the transgenic lines were not different to the rates observed for the WT when grown on MS agar medium for 3 d. However, the transgenic lines showed 20–32% and 6–13% higher seed germination than WT plants after mannitol (300 mM and 350 mM) and ABA (0.5 μM and 1 μM) treatments, respectively (Supplementary Fig. S2A, B). Given that ABA controls seed germination, and that its biosynthesis can be affected by abiotic stress (Iuchi *et al.*, 2001; Fujii and Zhu, 2009), we measured endogenous ABA contents. We found that ABA levels in the transgenic lines were slightly higher than in WT following mannitol or ABA treatments (Supplementary Fig. S2C).

We also examined cotyledon greening rates in plants grown on MS agar medium for 7 d and observed that the transgenic lines and WT showed no significant differences (Fig. 3A). However, the transgenic lines showed 30–39% and 19–20% higher cotyledon greening rates than WT plants after 7 d with mannitol (300 mM and 350 mM) and ABA (0.5 μM) treatments, respectively (Fig. 3B). The cotyledon greening rates of both transgenic lines and WT decreased substantially in response to the 1 μM ABA treatment, but the transgenic seedlings exhibited 4-6% higher cotyledon greening rates than the WT (Fig. 3A, B). The cotyledon greening rates were further tested at different time points after treatment with 350 mM mannitol or 1 μM ABA. As shown in Fig. 3C, neither the transgenic lines nor WT plants had greening cotyledons 3 d after treatment with 350 mM mannitol. However, the cotyledon greening rates of the transgenic lines were significantly higher than those of WT 4 d after treatment, and gap gradually increased with time, and peaked at 7 d. When grown on MS agar medium supplied with 1 μM ABA, the cotyledon greening rates of transgenic lines and WT showed no significant difference at 6, 7 and 8 d, while the transgenic lines exhibited 11-12% and 29-35% higher cotyledon greening rates than WT plants 9 d and 11 d after treatment, respectively (Fig. 3D). These results suggest that over-expressing *VlbZIP30* reduced sensitivity of *A. thaliana* seedlings to mannitol and ABA. In addition, the endogenous ABA levels in the transgenic lines were higher than those in WT after exposure to 350 mM mannitol (Fig. 3E), suggesting that the transgenic plants were insensitive to mannitol, possibly due to altered endogenous ABA levels.

**Fig. 3.**
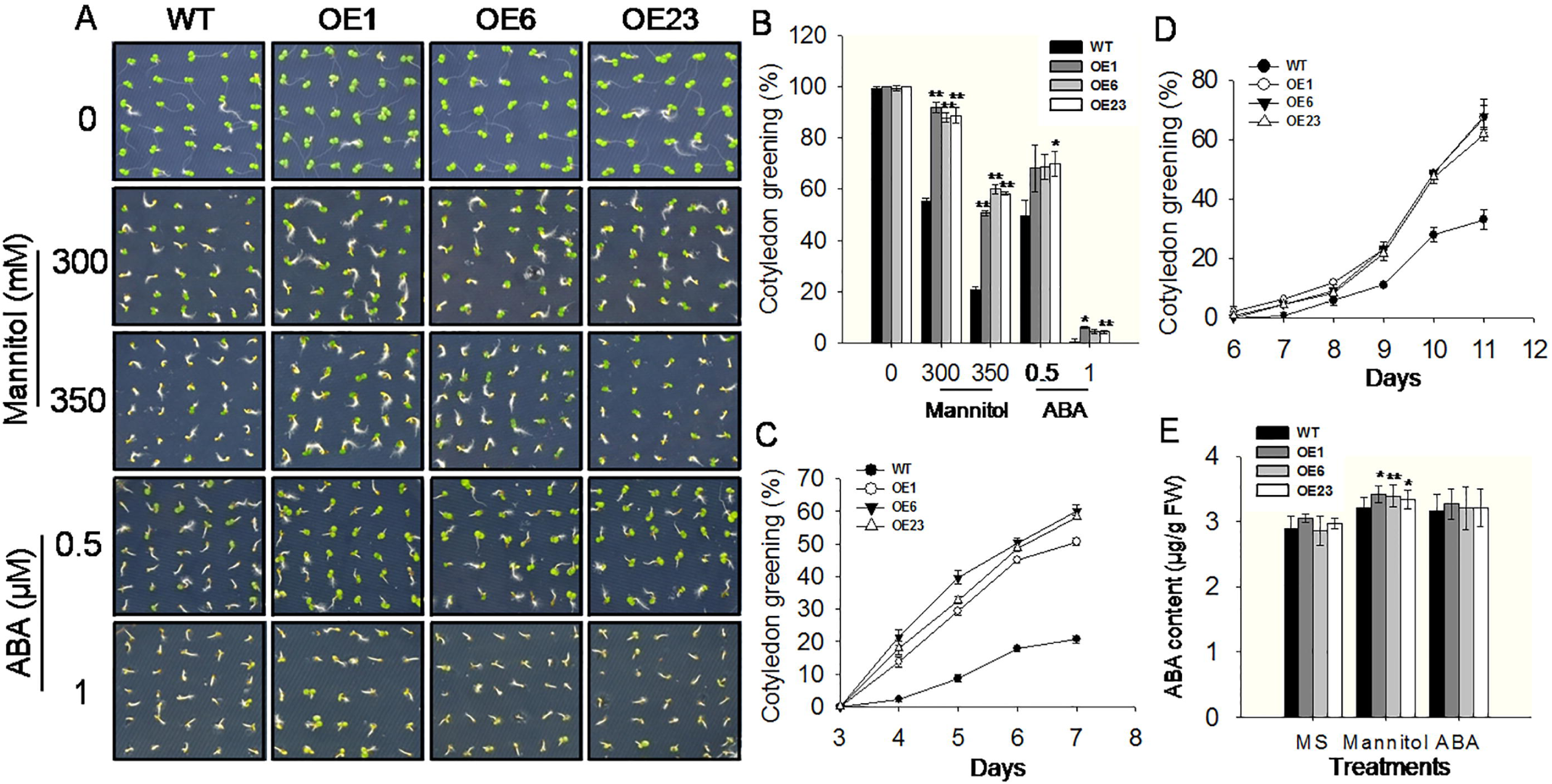
Phenotypes of wild type (WT) and *VlbZIP30* overexpressing (OE) transgenic lines at the greening cotyledon stage following mannitol and abscisic acid (ABA) treatments. (A) Greening cotyledons from WT and transgenic lines 7 d after seeds were cultivated on Murashige-Skoog (MS) agar medium, with or without 300 mM or 350 mM mannitol, or 0.5 μM or 1 μM ABA. (B) Cotyledon greening rates of WT and transgenic lines 7 d after cultivation on MS agar medium with or without 300 mM or 350 mM mannitol, or 0.5 μM or 1 μM ABA. (C) and (D) Cotyledon greening rates of WT and transgenic lines grown on MS basal medium containing 350 mM mannitol (C) or 1 μM ABA (D). (E) Endogenous ABA levels of WT and transgenic lines 7 d after cultivationon MS agar medium, or MS agar medium containing 350 mM mannitol or 1μM ABA. Three independent experiments were performed with ∼100 seeds per experiment. Error bars indicate ±SE. Asterisks indicate statistical significance (*0.01 < P < 0.05, **P < 0.01, Student’s *t*-test) between the transgenic and WT plants.

To further characterize morphological changes of WT and transgenic seedlings in response to the mannitol and ABA treatments, the various seed genotypes described above were grown on MS agar medium, with or without 300 mM mannitol, or 0.5 μM ABA for 14 d. As shown in Fig. 4A, the size of cotyledons and roots of the transgenic seedlings were significantly bigger and longer than those of WT following both treatments, while there was no differences observed under control conditions.

**Fig. 4.**
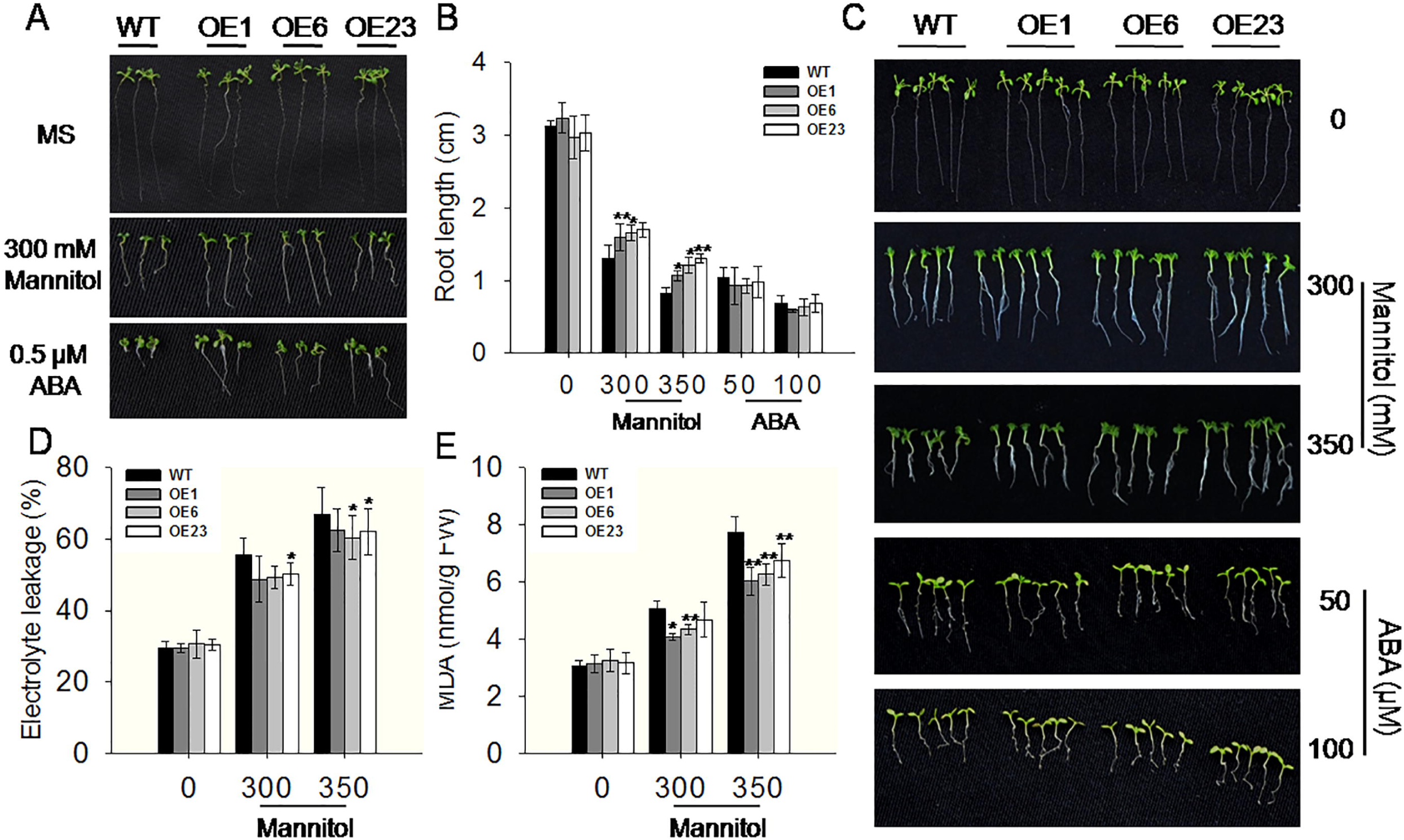
Phenotypes of wild type (WT) and *VlbZIP30*-overexpressing (OE) transgenic lines at the post-germination growth stage during mannitol and abscisic acid (ABA) treatments. (A) Photographs of morphology in WT and transgenic lines 14 d after seeds were cultivated on Murashige-Skoog (MS) agar medium with or without 300 mM mannitol or 0.5 μM ABA. (B) Root length of WT and transgenic lines after 7 d of growth with or without mannitol (300 mM or 350 mM) or ABA (50 μM or 100 μM). (C) Photographs of 14-day-old seedlings transferred from MS agar medium to MS agar medium or MS agar medium supplemented with mannitol (300 mM or 350 mM) or ABA (50 μM or 100 μM) for 7 d. Electrolyte leakage (D) and malondialdehyde (MDA) content (E) of 14-day-old seedlings transferred from MS agar medium to MS agar medium or MS agar medium supplemented with mannitol (300 mM or 350 mM) or ABA (50 μM or 100 μM) for 7 d. In all cases, data represent mean values ±SE from three independent experiments. Asterisks indicate statistical significance (*0.01 < P < 0.05, **P < 0.01, Student’s *t*-test) between the transgenic and WT plants.

To determine whether the transgenic lines had longer primary roots as a consequence of precocious germination or development of post-germination growth, 7-d-old transgenic lines and WT seedlings grown on MS agar medium were transferred to MS agar medium with or without mannitol or ABA, and grown for another 7 d. When grown on MS agar medium, the primary root lengths were similar; however, when grown on MS agar medium containing mannitol (300 mM and 350 mM) or ABA (50 μM and 100 μM), the transgenic lines had relatively longer primary roots than WT seedlings under mannitol treatment, while there was no significant difference in response to the ABA treatment (Fig. 4B, C). This indicated that *VlbZIP30* plays a role in suppressing the retardation of germination mediated by ABA, but not in root growth inhibition.

One effect of osmotic stress is membrane lipid peroxidation, and the levels of electrolyte leakage and malondialdehyde (MDA) are often used as indicators of the degree of cell membrane injury and tissue damage (Neill *et al.*, 2002; Pompelli *et al.*, 2010). We measured electrolyte leakage and MDA levels in the transgenic and WT seedlings after 300 mM or 350 mM mannitol treatment and saw that under normal growth conditions, there was no significant difference between the transgenic lines and WT seedlings. However, following treatments with different mannitol concentrations, the values in the transgenic lines were significantly lower than in WT, indicating that the degree of membrane and tissue damage was less as a result of *VlbZIP30-overexpression* (Fig. 4D, E). Taken together, these results indicated that overexpressing *VlbZIP30* in *A. thaliana* enhanced osmotic stress tolerance not only at the germination stage, but also during post-germination developmental processes.

### The expression of many ABA- or drought-responsive genes is induced in VlbZIP30-overexpressing lines

To examine the possible roles of *VlbZIP30* in transcriptional regulation in response to ABA and osmotic stress, we performed a global transcriptome analysis to identify DEGs between the WT and *VlbZIP30*-overexpressing lines using RNA-seq. Seeds from WT and OE1 transgenic lines were cultivated on MS agar medium with or without 0.5 μM ABA or 300 mM mannitol for 7 d, and the seedlings were then collected for transcriptome analysis (Fig. 3A, scheme summarized in Fig. 5A). DEGs were defined based on a threshold of 2-fold change (FDR<0.05). We identified 10 genes that were up- and 10 that were down-regulated in the OE lines compared with WT plants under control conditions (OEC / WTC, Fig. 5B, C). Details of these 20 genes, including their annotation and their expression levels are listed in Supplementary Table S2. After treatments, a total of 1,735 and 2,203 up-regulated genes and 1,734 and 1,764 down-regulated genes were identified in WT plants subjected to ABA (WTA / WTC) and mannitol (WTM / WTC) stress, respectively (Fig. 5B, C). A total of 1,510 and 1,494 up-regulated genes, and 1,058 and 729 down-regulated genes were found in the OE lines subjected to ABA (OEA / OEC) and mannitol (OEM / OEC) stress, respectively (Fig. 5B, C). We also identified 359 and 139 genes that were up- and down-regulated, respectively, in the OE lines compared with WT plants when treated with ABA (OEA / WTA, Fig. 5B, C), while 783 and 344 genes were up- and down-regulated, respectively, when treated with mannitol (OEM / WTM, Fig. 5B, C).

**Fig. 5.**
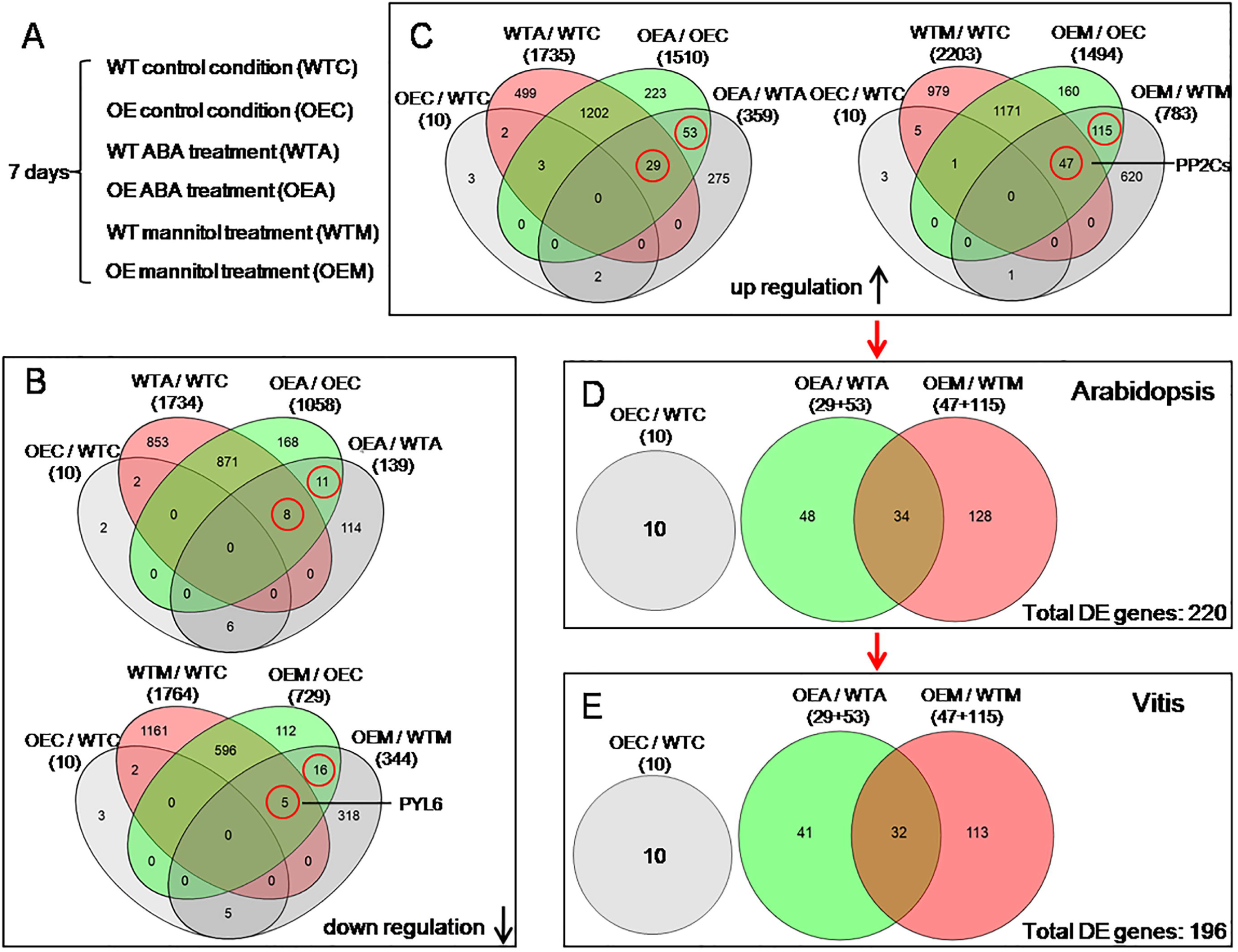
Venn diagram representation of the differentially expressed genes (DEGs) in four comparisons of wild type (WT) and *VlbZIP30-overexpressing* plants (OE) grown under control conditions, abscisic acid (ABA) or mannitol stress. (A) Experimental set up: WT and OE transgenic seeds were cultivated on Murashige-Skoog (MS) agar medium (WTC, OEC) or MS agar medium supplemented with 0.5 μM ABA (WTA, OEA) or 300 mM mannitol (WTM, OEM) for 7 d. (B) and (C) show the numbers of overlapping down-regulated and up-regulated genes, respectively. The numbers in brackets represent the total numbers of DEGs in different comparisons. The DEGs in red circles were selected for further analysis. (D) The selected up-regulated genes in the comparisons (OE / WT) under control conditions, ABA or mannitol stress in *Arabidopsis thaliana*. (E) The grapevine ( *Vitis vinifera*) homologs of the selected *A. thaliana* DEGs from (D).

To deduce the possible functions of the *VlbZIP30-induced* genes, we performed a gene ontology (GO) analysis of the genes, whose expression levels were significantly altered in the OE lines compared with the WT plants in response to ABA (OEA / WTA) or mannitol stress (OEM / WTM), using the PageMan profiling tool (Usadel *et al.*, 2006). This revealed that some genes encode TFs, and some genes are putatively involved in photosynthesis, stress, signaling, transport, development and several other metabolic pathways involving hormones, amino acids and lipids (Fig. 6).

**Fig. 6.**
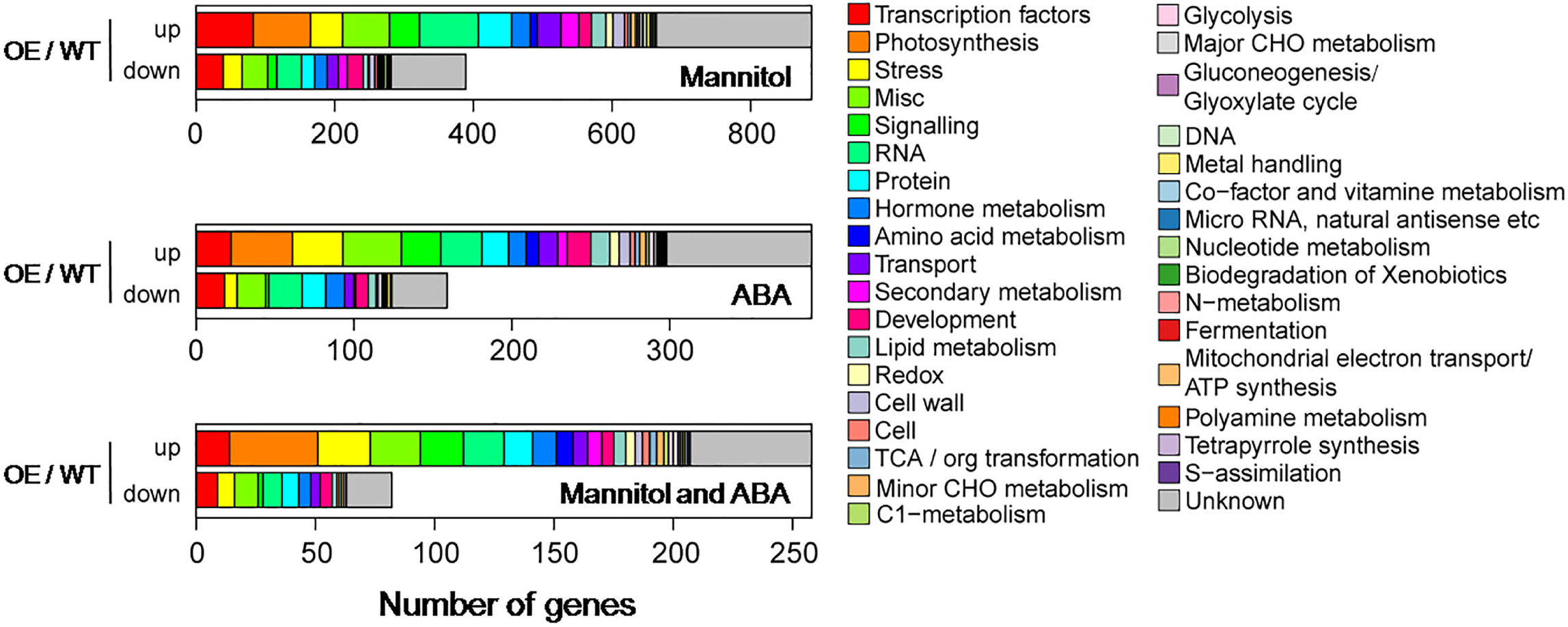
Gene ontology (GO) analyses of the differentially expressed genes (DEGs) in the *VlbZIP30-overexpressing* plants (OE) compared with wild type (WT) plants in response to abscisic acid (ABA) (OEA / WTA) or mannitol stress (OEM / WTM). Bar graphs separately display the numbers of DEGs classified into each GO category among the genes up- or down-regulated in the comparisons (OE / WT) induced by ABA, mannitol, or ABA and mannitol treatment. GO analyses are based on the PageMan profiling tool (Usadel *et al*, 2006) and the Arabidopsis Functional Modules Supporting Data (Heyndrickx and Vandepoele, 2012).

To better understand the role of *VlbZIP30* in ABA and osmotic stress signaling, the OEA / OEC and OEA / WTA intersecting genes (up-regulated genes, 29+53; down-regulated genes, 8+11) and the OEM / OEC and OEM / WTM intersecting genes (up-regulated genes, 47+115; down-regulated genes, 5+16) were selected for further analysis (Fig. 5B, C). We identified all of the 248 (210 up-regulated genes, 38 down-regulated genes) genes mentioned above, and found that 38% (95/248) genes had been identified in *A. thaliana.* Among them, 54 genes were involved in abiotic stress, including stress-responsive genes (*RD20, RD22, RD26, SIS* and *ERD10)*, ABA signaling genes (*AFP1, AFP3, HB7, HB12, NCED3, MAPKKK18, PYL6)*, PP2C genes (*ABI1, ABI2, HAI1, HAI2, HAB1* and *PP2CA)*, TFs (*ABF2, ABF3, ABF4*, *DREB1A, NFYA5, NFYB2, NAP, MYB74, WRKY28, ERF053* and *bHLH129)*, and others (Table 1). The other 41 genes were involved in other processes such as wax biosynthesis (*ABCG19*), photosynthesis, transport, hormone signaling (brassinosteroids, ethylene and cytokinins), mineral homeostasis (Fe, Ca, Pi and S) and others (Supplementary Table S3). We noted that the genes involved in hormone signaling were almost all down-regulated in the OE lines compared with WT plants in response to both treatments, while the genes involved in mineral homeostasis were almost all up-regulated in the OE lines compared with WT plants following ABA treatment (Supplementary Table S3). To date, the functions of these genes characterized in Arabidopsis were consistent with the results of GO analysis (Fig. 6).

**Table 1.** Selected genes involved in abiotic stress with expression changes (FDR<0.05) of at least twofold in the *VlbZIP30* transgenic plants under different experiments from the microarray.

| Locus | Symbol | Description | Reference | Log<sub>2</sub>FC (OE / WT) |  |  |
| --- | --- | --- | --- | --- | --- | --- |
|  |  |  |  | C | M | A |
| Stress-responsive gene |  |  |  |  |  |  |
| AT2G33380 | RD20 | stress-inducible caleosin | Aubert *et al.*, 2010 | -0.421 | 2.781 | 2.083 |
| AT5G25610 | RD22 | responsive to dehydration protein | Yoshida *et al.*, 2002 | -0.080 | 1.160 | 0.121 |
| AT4G27410 | RD26 | dehydration-induced NAC protein | Fujita *et al.*, 2004 | 0.187 | 1.325 | 0.560 |
| AT2G42530 | COR15B | cold-regulated protein | Nelson *et al.*, 2007 | 0.329 | 1.833 | 1.511 |
| AT2G18050 | HIS1-3 | dehydration-inducible histone gene | Zhang *et al.*, 2008 | 0.711 | 1.106 | -0.089 |
| AT5G02020 | SIS | involved in salt tolerance | Brinker *et al.*, 2010 | 0.176 | 1.530 | 1.219 |
| AT1G20450 | ERD10 | early responsive to dehydration | Kovacs *et al.*, 2008 | 0.217 | 0.906 | 1.214 |
| ABA signaling |  |  |  |  |  |  |
| AT1G69260 | AFP1 | ABI five binding protein | Garcia *et al.*, 2008 | 0.187 | 1.275 | 0.646 |
| AT3G29575 | AFP3 | ABI five binding protein | Garcia *et al.*, 2008 | 0.286 | 1.256 | 0.560 |
| AT4G26080 | ABI1 | protein phosphatase 2C protein | Merlot *et al.*, 2001 | 0.186 | 1.217 | -0.060 |
| AT5G57050 | ABI2 | protein phosphatase 2C protein | Merlot *et al.*, 2001 | 0.147 | 1.192 | 0.303 |
| AT5G59220 | HAI1 | protein phosphatases 2C protein | Bhaskara *et al.*, 2012 | 0.219 | 2.543 | 1.931 |
| AT1G07430 | HAI2 | protein phosphatases 2C protein | Bhaskara *et al.*, 2012 | 0.887 | 2.247 | 0.365 |
| AT1G72770 | HAB1 | protein phosphatases 2C protein | Saez *et al.*, 2006 | -0.003 | 1.255 | 0.675 |
| AT3G11410 | PP2CA | protein phosphatases 2C protein | Yoshida *et al.*, 2006 | 0.384 | 1.634 | -0.077 |
| AT2G46680 | HB7 | involved in ABA signaling pathway | Valdes *et al.*, 2012 | -0.017 | 1.555 | 0.619 |
| AT3G61890 | HB12 | involved in ABA signaling pathway | Valdes *et al.*, 2012 | -0.029 | 2.007 | 0.879 |
| AT3G14440 | NCED3 | key enzyme in ABA biosynthesis | Iuchi *et al.*, 2001 | 0.833 | 1.845 | 0.125 |
| AT1G05100 | MAPKKK18 | involved in ABA signaling pathway | Tajdel *et al.*, 2016 | ND | 1.550 | 1.137 |
| AT2G40330 | PYL6 | core regulatory component of ABA receptor | Gonzalez-Guzman *et al.*, 2012 | -0.342 | -1.213 | ND |
| Transcription factor |  |  |  |  |  |  |
| AT4G05100 | MYB74 | R2R3-MYB family gene | Xu *et al.*, 2015 | 0.275 | 1.672 | 0.691 |
| AT4G21440 | MYB102 | R2R3-MYB family gene | Denekamp and Smeekens, 2003 | ND | 1.630 | 0.332 |
| AT4G34000 | ABF3 | ABRE-binding transcription factor | Kang *et al.*, 2002 | 0.013 | 1.129 | 0.565 |
| AT1G45249 | ABF2 | ABRE-binding transcription factor | Fujita *et al.*, 2005 | 0.001 | 1.954 | 0.722 |
| AT3G19290 | ABF4 | ABRE-binding transcription factor | Kang *et al.*, 2002 | 0.053 | 1.050 | 0.473 |
| AT4G25480 | DREB1A | DRE-binding transcription factor | Wei *et al.*, 2016 | 0.286 | 2.020 | -0.402 |
| AT1G54160 | NFYA5 | CCAAT-binding transcription factor | Li *et al.*, 2008 | 0.342 | 1.469 | 0.367 |
| AT5G47640 | NFYB2 | His-like transcription factor | Kumimoto *et al.*, 2013 | 0.200 | 2.885 | 0.755 |
| AT1G69490 | NAP | NAC family transcription factor | Sakuraba *et al.*, 2015 | 0.377 | 1.297 | 0.867 |
| AT4G09820 | TT8 | bHLH transcription factor | Rai *et al.*, 2016 | -0.591 | 1.963 | 0.201 |
| AT4G18170 | WRKY28 | WRKY transcription factor | Babitha *et al.*, 2013 | -0.577 | 1.277 | 0.594 |
| AT2G20880 | ERF053 | ethylene response factor | Hsieh *et al.*, 2013 | ND | ND | 3.986 |
| AT2G43140 | BHLH129 | bHLH transcription factor | Tian *et al.*, 2015 | -0.21 | -1.005 | ND |
| AT3G23250 | MYB15 | R2R3-MYB family gene | Li *et al.*, 2010 | -0.098 | -1.527 | -0.437 |
| Disease |  |  |  |  |  |  |
| AT5G59310 | LTP4 | susceptibility to pseudomonas | Gao *et al.*, 2016 | 0.198 | 1.467 | 0.086 |
| AT1G75040 | PR5 | pathogenesis-related gene | Liu *et al.*, 2013 | -0.542 | 2.521 | ND |
| AT1G54040 | ESP | involved in necrotrophic fungi resistance | Buxdorf *et al.*, 2013 | ND | 2.064 | 1.439 |
| **Hsp** |  |  |  |  |  |  |
| AT3G22830 | HSFA6B | heat stress transcription factor | Huang *et al.*, 2016 | 0.129 | 1.648 | 0.465 |
| AT5G43840 | HSFA6A | heat stress transcription factor | Hwang *et al.*, 2014 | ND | 2.398 | 1.028 |
| **Metabolism** |  |  |  |  |  |  |
| AT2G39800 | P5CS1 | involved in proline biosynthesis | Martinez *et al.*, 2015 | 0.251 | 1.798 | 1.564 |
| AT1G56650 | MYB75 | involved in anthocyanin metabolism | Lee *et al.*, 2016b | ND | 1.730 | 0.606 |
| AT3G63060 | EDL3 | regulate anthocyanin accumulation | Koops *et al.*, 2011 | -0.121 | 1.232 | 0.026 |
| AT2G05100 | LHCB2.1 | chlorophyll a/b-binding proteins | Lucinski and Jackowski, 2013 | -0.311 | 1.240 | 0.651 |
| AT2G05070 | LHCB2.2 | chlorophyll a/b-binding proteins | Lucinski and Jackowski, 2013 | -0.294 | 1.601 | 0.818 |
| AT3G27690 | LHCB2.4 | chlorophyll a/b-binding proteins | Lucinski and Jackowski, 2013 | -0.103 | 1.920 | 1.153 |
| AT3G47600 | MYB94 | activate cuticular wax biosynthesis | Lee *et al.*, 2016a | 0.134 | 1.913 | 0.312 |
| AT5G62470 | MYB96 | activate cuticular wax biosynthesis | Lee *et al.*, 2016a | 0.241 | 2.048 | 0.738 |
| AT2G33790 | ATAGP30 | arabinogalactan protein | van Hengel and Roberts, 2003 | 0.543 | -0.042 | 1.253 |
| **Transporter** |  |  |  |  |  |  |
| AT1G22370 | AtUGT85A5 | UDP-glycosyl transferase | Sun *et al.*, 2013 | 0.051 | 1.118 | 0.621 |
| AT4G13420 | HAK5 | high affinity K<sup>+</sup> transporter | Brauer *et al.*, 2016 | 0.460 | 0.154 | -1.043 |
| AT2G02930 | ATGSTF3 | glutathione transferase (GST) | Lee *et al.*, 2014 | -0.316 | -0.977 | -1.478 |
| **Enzyme** |  |  |  |  |  |  |
| AT1G56600 | AtGolS2 | galactinol synthase | Selvaraj *et al.*, 2017 | -0.059 | 1.157 | -0.073 |
| AT1G60190 | PUB19 | U-Box E3 Ubiquitin Ligase | Liu *et al.*, 2011b | -0.059 | 1.157 | -0.073 |
| AT1G69270 | RPK1 | leucine-rich receptor-like kinase | Osakabe *et al.*, 2010 | 0.148 | 1.057 | -0.006 |

<-1 -1~0 0~1 1~2 >2 ND
FDR, false discovery rate; FC, fold change; OE / WT, overexpression line / wild type; M, mannitol; A, ABA; ND,
not detected

To confirm the RNA-seq results, we examined the expression of 20 drought-responsive genes by qRT-PCR and saw that the expression changes of all these genes were similar in the RNA-seq and qRT-PCR data. As shown in Fig. 7, the expression of four stress-marker genes and six ABA signaling genes, but not *NCED3*, was up-regulated during both treatments in OE lines compared with WT plants. There were increased transcript levels of the ABA biosynthesis related gene *NCED3* in the OE lines during mannitol stress, which correlated with the endogenous ABA content (Fig. 3E). Furthermore, the expression of several marker genes in the core ABA signaling network, including PP2Cs and *PYL6*, also showed significant change. Specifically, the expression of 4 PP2Cs significantly increased in the OE lines relative to WT plants, while the expression of *PYL6* decreased in OE lines compared with WT plants during mannitol stress.

**Fig. 7.**
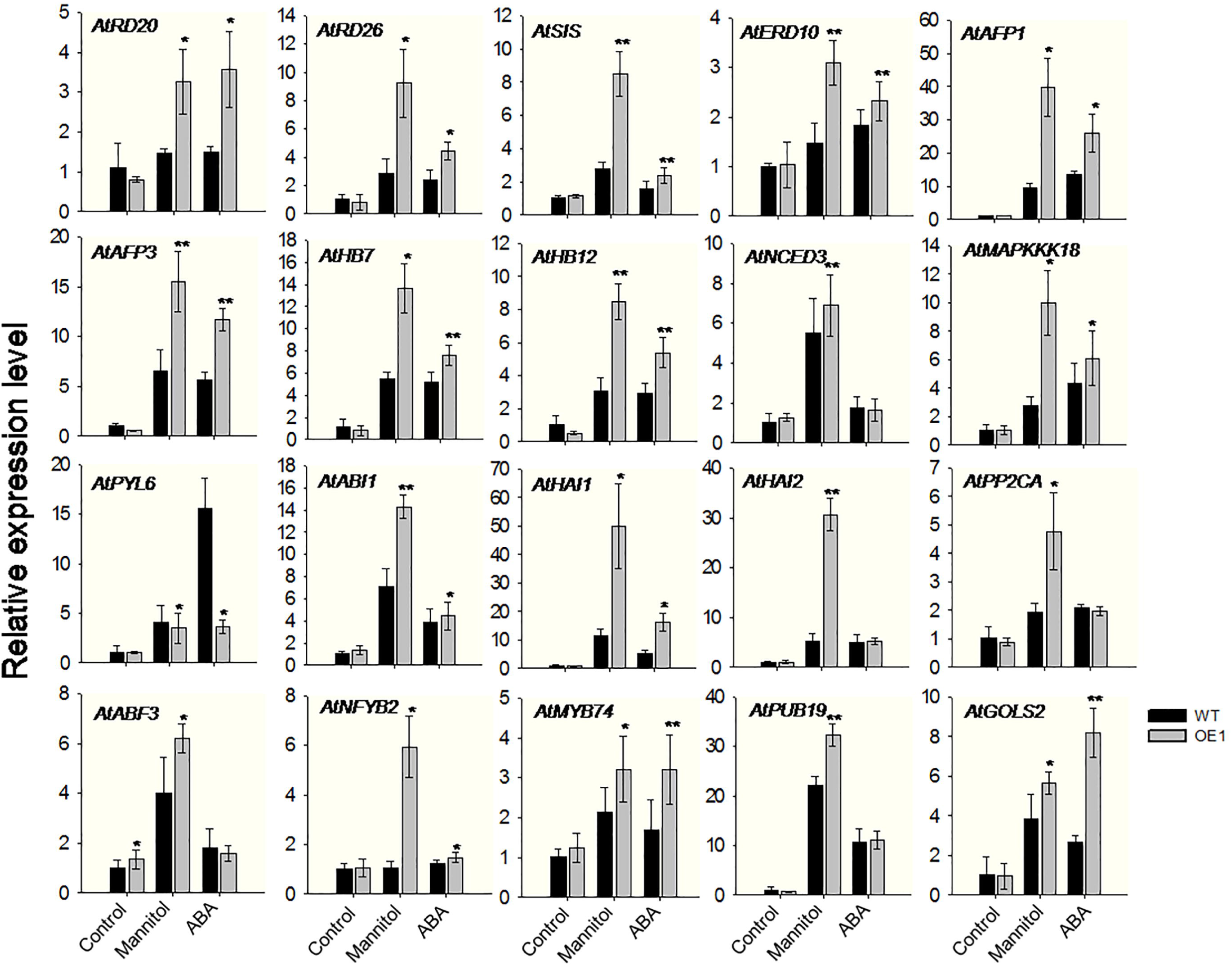
Experimental validation of transcriptome data by expression analysis of representative drought-responsive genes in *Arabidopsis thaliana* using qRT-PCR. The expression level in the wild type (WT) plants under control conditions was defined as 1. 0. The *AtActin2* gene was used as an internal control. Data represent mean values ±SE from three independent experiments. Asterisks indicate statistical significance (*0.01 < P < 0.05, **P < 0.01, Student’s *t*-test) between the overexpressing (OE) lines and WT plants.

### The presence of a potential G-box motif in VlbZIP30 induced genes

To identify candidate *VlbZIP30* target genes, promoter analyses of the DEGs between OE lines and WT plants under control conditions (10 up-regulated and 10 down-regulated genes) and during ABA or mannitol treatment (210 up-regulated and 38 down-regulated genes) were performed using the DREME motif discovery tool. All up-regulated and all down-regulated genes were divided in two clusters performed promoter analyses, respectively. Excitingly, We identified a potential G-box *cis*-element motif (ACGTGKV; E-value, 2.5e-017) including a 4-bp core sequence (ACGT) that is known to be a bZIP binding motif, as being significantly enriched in the promoters of the up-regulated genes (Fig. 8A), in that 85% (187/220) carried this motif in their upstream 1,500 bp promoter region. The G-box motif was not enriched in the down-regulated genes. We next analyzed the number and location of the G-box motifs in the 187 gene promoters, and observed that many genes carried 1 to 3 G-box motifs, and that the highest G-box frequency was within the first 300 bp from the start codon site (Fig. 8C, E).

**Fig. 8.**
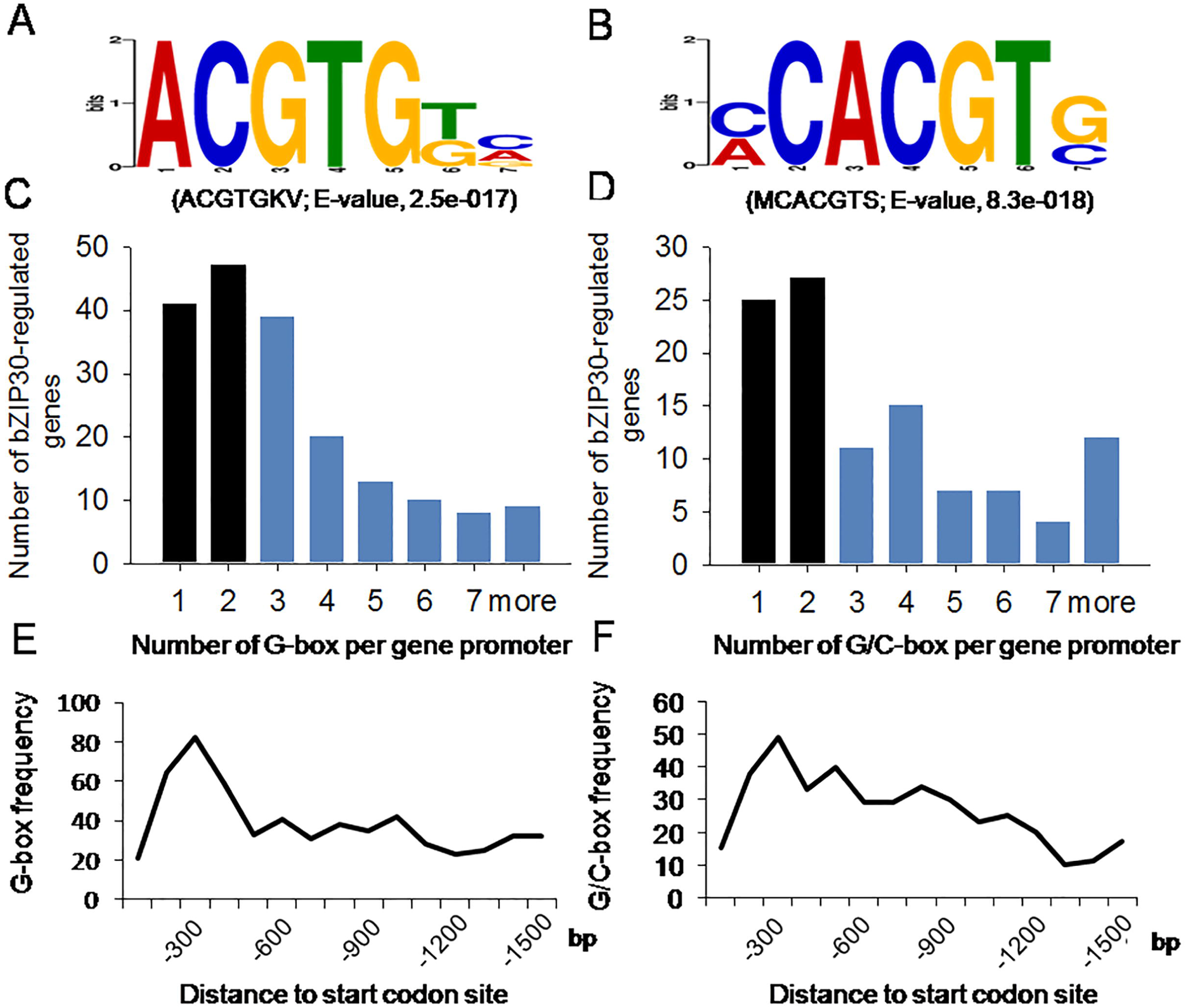
Enrichment of a potential G-box motif in *VlbZIP30* induced genes. (A) and (B) DREME motif analysis, showing the predicted G-box (ACGTGKV) motif (A) in the promoter regions of selected *Arabidopsis thaliana* up-regulated genes from the overexpressing (OE) lines, and the potential G/C-box (MCACGTS) motif (B) in the promoter regions of the corresponding grapevine (*Vitis vinifera*) homologs. (C) and (D) Number of predicted G-box motifs in the promoters of the selected *A. thaliana* genes (C), and the number of predicted G/C-box motifs in the promoter of the corresponding grapevine homologs (D). Predicted *VlbZIP30-induced A. thaliana* and grapevine genes with at least three G-box or G/C-box motifs were selected for further analyses (highlighted in blue). (E) and (F) Frequency of the predicted G-box motif in the promoters of the selected *A. thaliana* genes with respect to their distance from the start codon site (E), and the frequency of the potential G/C-box motif in the promoters of the corresponding grapevine homologs with respect to their distance from the start codon site (F).

ABRE (PyACGTGGC) and G-box (CACGTG) elements have previously been identified as *cis*-binding elements for bZIP proteins that regulate gene expression in response to ABA or drought stress in many plants, such as *A. thaliana* (Uno *et al.*, 2000), rice (Liu *et al.*, 2014), wheat (Wang *et al.*, 2016), soybean (Liao *et al.*, 2008), tomato (Hsieh *et al.*, 2010), maize (Zhang *et al.*, 2011), orange (Zhang *et al.*, 2015), potato (Muniz Garcia et al., 2012), and *Tamarixhispida* (Ji *et al.*, 2015). We searched the grapevine genome for homologs of the identified *A. thaliana* up-regulated (220) and down-regulated (48) genes (Fig. 5D, E; Supplementary Data S1) and a promoter analysis performed using the DREME motif discovery tool. Homologs were found of 195 of the up-regulated and 44 of the down-regulated genes (Supplementary Data S1). Using the same analytical method, surprising, a potential G/C-box *cis*-element motif (MCACGTS; E-value, 8.3e-018) including the core sequence (ACGT) was found to be significantly enriched in the homologs to the up-regulated genes (Fig. 8B). We determined that 55% (108/195) of the up-regulated genes had at least one G/C-box motif in their upstream 1,500 bp promoter region, while this motif was not enriched in the down-regulated genes. Many of the 195 genes carried 1 or 2 G/C box motifs, and the highest frequency was seen in these quence up to 300 bp from start codon site (Fig. 8D, F), similar to the *A. thaliana* genes (Fig. 8C, E).

Next, those genes with three or more G-box (A. *thaliana*) or G/C-box (grapevine) motifs were selected (Piya *et al.*, 2017). We found that 52% (97/187) of the *A. thaliana* genes and 50% (54/108) of the grapevine genes had at least three motifs and that all of the latter had G-box, but not C-box motifs. We scanned the above-mentioned 97 *A. thaliana* genes and 54 grapevine genes containing three or more G-box motifs, and found that 61 of the *A. thaliana* genes were homologous to one of the 54 grapevine genes. Of the 97 and 61 *A. thaliana* genes, we identified 39 with three or more G-box motifs in their promoters by Venn diagram (Fig. 9A). Another DREME promoter analysis of these 39 genes revealed 4 enriched motifs, including a perfect G-box (CACGTG, E-value: 9.0e-012), a GAGA-box (DAGAGAGA, E-value: 1.1e-005), an AAGAAAAR motif (E-value: 7.9e-004), and a TATA-box (ABATATAT, E-value: 9.9e-004) (Fig. 9B). The frequencies of the G-box in the *VlbZIP30-induced* 39 and 220 *A. thaliana* genes were 89.7% and 70.0%, respectively, while the frequency in the whole genome was 51.5%, suggesting an enrichment in the *VlbZIP30* induced genes. Such an enrichment was not found for the other three motifs (Fig. 9B).

**Fig. 9.**
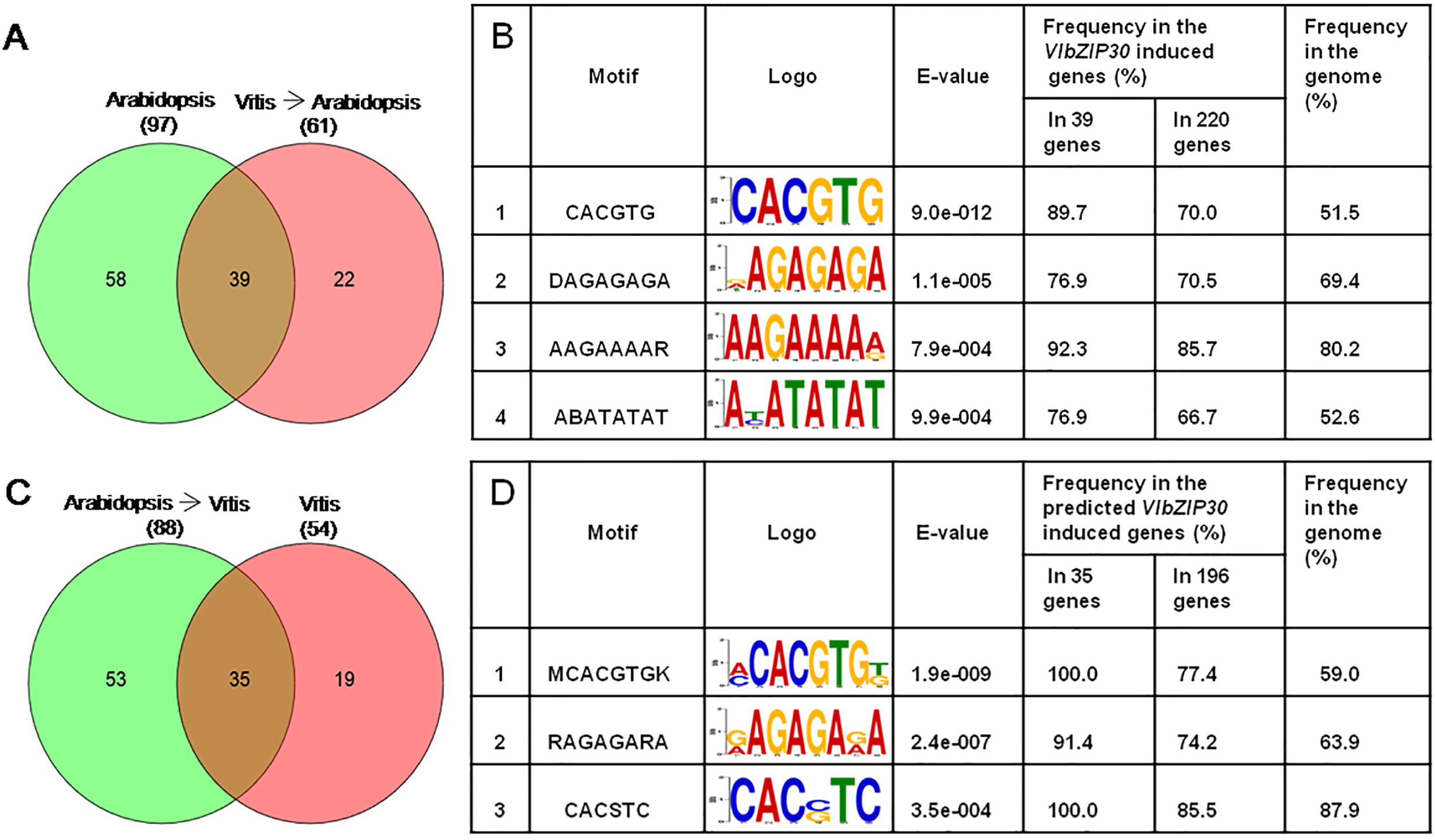
*In silico* promoter analyses of the selected *Arabidopsis thaliana* genes and the grapevine (*Vitis vinifera*) homologs. (A) Venn diagram showing the selected *A. thaliana* genes with 3 or more predicted G-box elements and the *A. thaliana* homologs of selected grapevine genes with 3 or more predicted G-box motifs. (C) Venn diagrams of the selected grapevine genes with 3 or more predicted G-box motifs and the grapevine homologs of the selected *A. thaliana* genes with 3 or more predicted G-box motifs. (B) and (D) The 1,500 bp promoter regions of the overlapping 39 *A. thaliana* genes from (A) and 35 grapevine genes from (C) were analyzed using the DREME motif enrichment tool. The potential *A. thaliana* G-box and grapevine G-box motif enrichments were identified using the whole genome sequences from *A. thaliana* and grapevine as a reference.

In grapevine, 88 homologs of the 97 *A. thaliana* genes were found. Among the 88 and 54 grapevine genes, we identified 35 with at least three G-box motifs in their promoters by Venn diagram (Fig. 9C). Three enriched motifs, including a potential grapevine G-box (MCACGTGK, E-value: 1.9e-009), a GAGA-box (RAGAGARA, E-value: 2.4e-007), and a CACSTC (E-value: 3.5e-004) motif were identified in these 35 genes (Fig. 9D). Among them, the frequencies of the G-box in the predicted *VlbZIP30-induced* 35 and 196 grapevine genes were 100.0% and 77.4%, respectively, while the frequency of the G-box in the whole genome was 59.0%, again indicating an enrichment in the predicted *VlbZIP30* induced genes. These results suggest that the 35 grapevine genes may be regulated by *VlbZIP30* via the potential G-box. The names and gene IDs of the 35 grapevine genes and 39 *A. thaliana* genes are listed in Supplementary Data S2.

### The expression of the predicted VlbZIP30 induced genes with at least three potential G-box motifs was up-regulated in grapevine following both ABA and drought treatments

To investigate the potential roles of the 35 identified grapevine genes in ABA and drought stress, two different grapevine-related databases for ABA (Pilati *et al.*, 2017) and drought (Rocheta *et al.*, 2016) stress treatments were analyzed. RNA-seq analysis was performed of grapevine berry skins with or without ABA treatment for 20 h and 44 h. A GrapeGene GeneChips^®^ data analysis was performed of leaves from two *Vitis vinifera* L. varieties (Trincadeira, TR and Touriga Nacional, TN) grown under control and drought greenhouse conditions, as well as fully irrigated and non-irrigated field conditions. We compared the genes in those datasets with the 35 predicted grapevine genes, and found that 31 and 25 appeared in the RNA-seq and GeneChips^®^ data (Fig. 10A). Of these, 74% (23/31) and 84% (21/25) were significantly up-regulated following the ABA and drought treatments, respectively (Fig. 10A). Six genes only responded to ABA and 4 only responded to drought, and 17 responded to both treatments (Fig. 10A). The detailed expression data for these 27 genes (Fig. 10A) are shown in heat map diagrams in Fig. 10B.

**Fig. 10.**
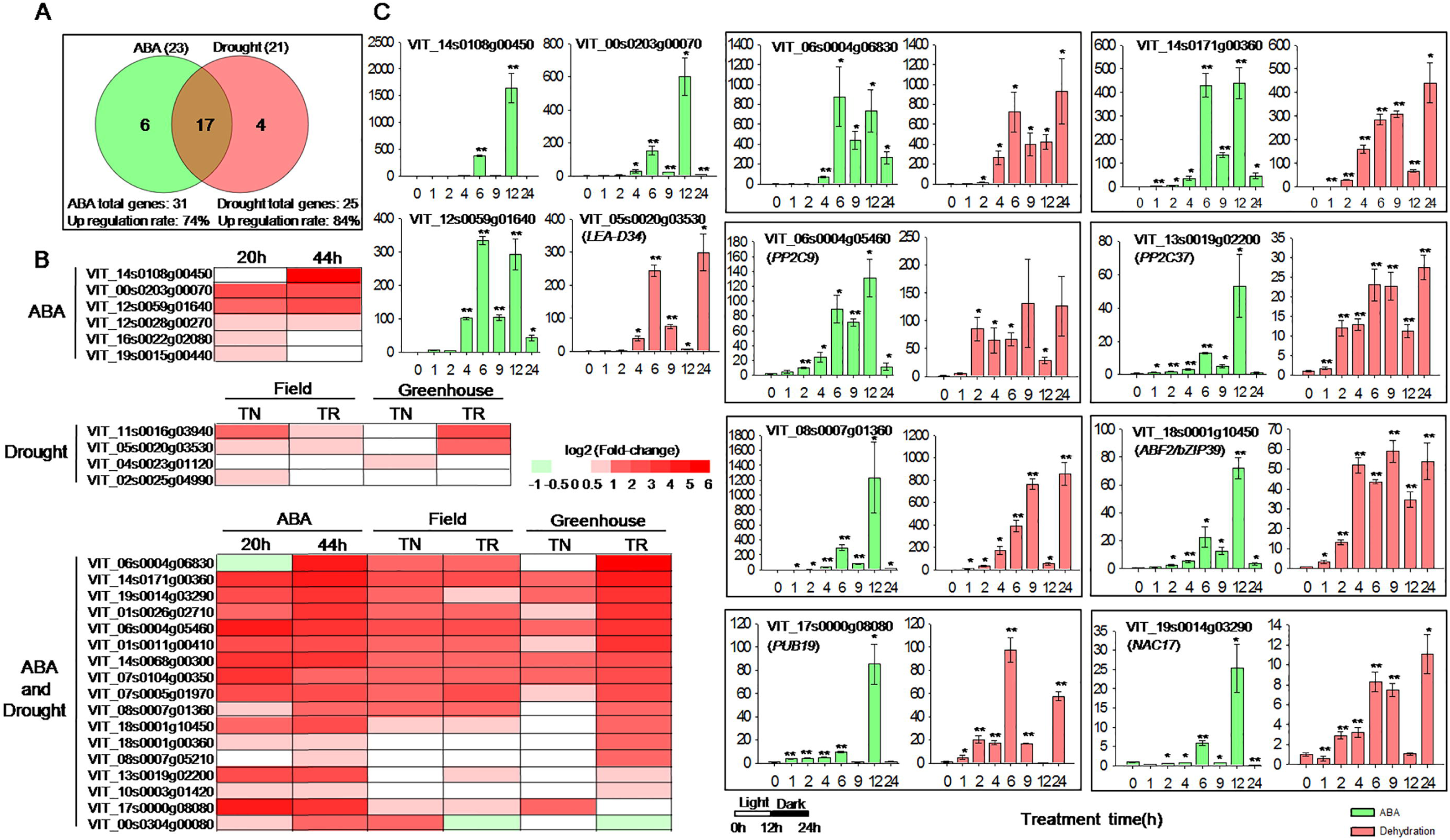
The expression profiles of predicted *VlbZIP30* induced genes in grapevine ( *Vitis vinifera*) following abscisic acid (ABA) or drought treatment. (A) Venn diagram of the selected up-regulated genes in two different grapevine-related databases (for ABA; Pilati *et al.*, 2017 and for drought stress; Rocheta *et al.*, 2016) with the 35 predicted grapevine genes. Thirty-one and 25 predicted grapevine genes were present in the RNA-seq and GeneChips^®^ data. The numbers in brackets represent the numbers of up-regulated genes in the two different grapevine-related databases. (B) Heat maps showing the expression of the 27 genes found in (A). RNA-seq analysis was performed of transcripts expressed in the skin of berries from grapevines treated, or not, with ABA for 20 h and 44 h. The GrapeGene GeneChips^®^ data was derived from an analysis of the leaves of two *V. vinifera* L. varieties (Trincadeira, TR and Touriga Nacional, TN) grown under control and drought conditions in a greenhouse, as well as under fully irrigated and non-irrigated conditions in the field. Heat map color gradation in red indicates the increase in expression (log2 fold change). (C) Gene expression profiles of randomly selected *VlbZIP30*-induced grapevine candidate genes analyzed using qRT-PCR. For each gene, the expression level in the 0 h sample from the ABA and dehydration treatments was defined as 1.0. The *VvActin1* gene was used as an internal control. Data represent mean values ±SE from three independent experiments. Asterisks indicate statistical significance (*0.01 < P < 0.05, **P < 0.01, Student’s *t*-test) between the treated and untreated control plants.

We randomly selected 16 genes of the 27 genes induced by ABA or drought stress, for confirmatory qRT-PCR expression analysis, using the previously mentioned grapevine leaf samples subjected to ABA or dehydration treatment. The results were consistent with the results previously published (Fig. 10C; Supplementary Fig. S3). Interestingly, *ABF2/bZIP39* (VIT_18s0001g10450), which was characterized as being involved in ABA signaling in grapevine cell culture, has been reported to transiently trans-activate the expression of *NAC17* (VIT_19s0014g03290) and *PUB19* (VIT 17s0000g08080) following ABA treatment (Nicolas *et al.*, 2014; Pilati *et al.*, 2017). In addition, overexpression of this gene in *A. thaliana* enhances tolerance to drought stress through the ABA signaling pathway (Tu *et al.*, 2016a), suggesting that *NAC17* and *PUB19* may enhance drought stress in grapevine.

### VlbZIP30 overexpressing A. thaliana lines showed enhanced dehydration tolerance at the adult stage

To further investigate the potential function of *VlbZIP30* in dehydration stress, the *VlbZIP30-overexpressing* (OE1, OE6 and OE23) lines and WT plants were grown for 3 weeks under normal growth conditions and then exposed to dehydration stress by withholding water for 8 d. All of the WT plants showed severe wilting symptoms, while only slight wilting was observed in the OE lines (Fig. 11A). Only 25% of the WT plants recovered after 3 d of rehydration, while the OE lines rapidly recovered, and 69-94% of the OE lines survived (Fig. 11A). We excised the aerial parts of the OE lines and WT plants and water loss was examined over time. The OE lines lost water more slowly than WT plants (Fig. 11B, C) and, consistent with the visible phenotypes, when leaves were stained with trypan blue, those of WT showed a deeper staining than those of the OE lines (Fig. 11B), suggesting a higher rate of cell death after 3 h dehydration. Since ROS triggered by drought stress can cause oxidative damage to cellular membranes, and ultimately result in cell death (You *et al.*, 2014), we measured ROS levels, as well as the activities of antioxidant enzymes (SOD, CAT and POD) in WT and OE lines before and after dehydration for 3 h. WT plants accumulated more ROS and had lower antioxidant enzyme activities than OE lines after 3 h dehydration (Supplementary Fig. S4), consistent with the cell death data. These results suggest that *VlbZIP30* promotes dehydration stress responses.

**Fig. 11.**
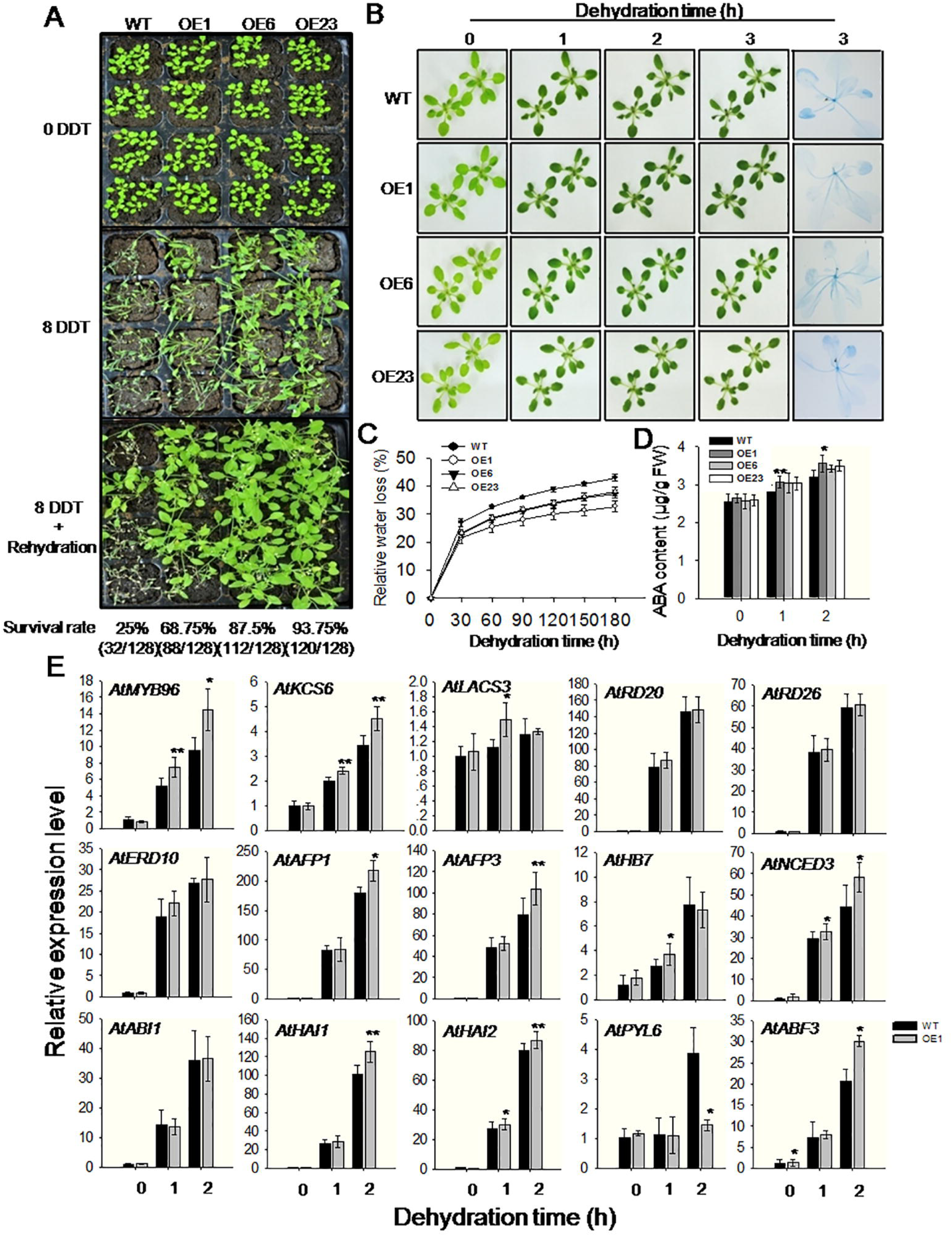
*VlbZIP30-overexpressing Arabidopsis thaliana* show enhanced dehydration tolerance at the adult stage. (A) Drought tolerance phenotypes and survival rates of wild type (WT) and transgenic (OE) lines grown in soil. Three-week-old plants (upper panel) were dehydrated for 8 d (middle panel) and then rehydrated for 3 d (lower panel). DDT: day of drought treatment. (B) Leaf phenotype of 3-week-old WT and OE lines before and after dehydration for 1, 2, 3 h. Staining with trypan blue of leaves detached from WT and OE lines and left to dehydrate for 3 h. (C) Relative water loss rates in WT and OE lines during 3 h of dehydration. (D) Endogenous abscisic acid (ABA) levels of WT and OE lines before and after dehydration for 1 h and 2 h. (E) Expression profiles of stress-related genes in WT and OE analyzed using qRT-PCR. Expression levels were based on total RNA extracted from the aerial parts of 3-week-old WT and OE lines that had been dehydrated on dry filter paper for 0, 1, 2 h. The *AtActin2* gene was used as an internal control. In all cases, data represent mean values ±SE from three independent experiments. Asterisks indicate statistical significance (*0.01 < P < 0.05, **P < 0.01, Student’s *t*-test) between the OE and WT plants.

It has been reported that some guard cells of drought-tolerant plants are hypersensitive to ABA (Fujita *et al.*, 2005; Sakuraba *et al.*, 2015), and this prompted us to test whether the enhanced dehydration resistance of the OE lines was associated with ABA-regulated stomatal closure. We measured stomatal aperture in the leaves of 3-week-old WT and OE lines in the presence of 10 μM ABA, but after 1 h of ABA treatment, no significant difference was observed (Supplementary Fig. S5). We then measured the endogenous ABA content before and after dehydration and while there was no difference under normal conditions, the ABA content of the OE lines was higher than in WT plants after dehydration for 1 h or 2 h (Fig. 11D). These results suggested that the OE lines had enhanced drought resistance due to ABA signaling, but not because of ABA-regulated stomatal closure.

Subsequently, by reading a large number of articles and analyzing the transcriptome data, we identified 6 genes (*MYB94, MYB96, KCS6, KCS12, LACS3*, and *ABCG19*) involved in cuticular wax biosynthesis that were significantly up-regulated in the OE lines compared with WT plants under mannitol stress (Table 1; Supplementary Table S3). Previous studies showed that *A. thaliana* can adapt to drought stress through *MYB94-* and MYB96-mediated regulation of cuticular wax biosynthesis via a downstream branch of the ABA core signaling pathway that is different from the stomata closure sub-branch of ABA signaling (Seo *et al.*, 2011; Cui *et al.*, 2016; Lee *et al.*, 2016a). Seo *et al.* (2011) suggested that the *MYB96* TF promotes drought resistance by regulating cuticular wax biosynthetic genes (including *KCS6, KCS12, LACS3*, and *ABCG19*) in the ABA-dependent pathway. We found that the expression of *MYB96* gradually increased after 1 and 2 h of dehydration stress in both WT and OE lines, but that it significantly increased in the OE lines compared with the WT (Fig. 11E). The expression level of *KCS6*, a *MYB96* direct binding target (Seo *et al.*, 2011; Lee *et al.*, 2016a), showed the same trend. The expression of another cuticular wax biosynthetic gene, *LACS3*, was also up-regulated in the OE lines compared with WT after dehydration for 1 h. However, the expression levels of the stress-marker genes, *RD20, RD26* and *ERD10*, were not significantly different between WT and OE lines in response to dehydration. *AFP1* and *AFP3*, which encode ABI five binding proteins, regulate the drought response in germinating *A. thaliana* seeds and seedlings, and their mutation results in ABA hypersensitivity (Garcia *et al.*, 2008). We observed that the expression of these genes significantly increased in OE lines compared with WT after 2 h of dehydration stress. The expression of the ABA core signaling network genes *ABI1, HAI1, HAI2* (PP2Cs), and *PYL6* was also measured, and while *HAI1* and *HAI2* were significantly up-regulated in the OE lines after 2 h of dehydration stress, the expression of *PYL6* was significantly down-regulated. *HB7* and *HB12*, which are positive transcriptional regulators of PP2Cs, suppress the transcription of *PYL5* and *PYL8* in response to ABA, and enhance drought tolerance as mediators of a negative feedback effect on ABA signaling in *A. thaliana* (Valdes *et al.*, 2012). Here, we found that the expression of *HB7* was up-regulated in the OE lines after 1 h of dehydration stress compared with the WT. In addition, transcript levels of the ABA biosynthesis marker gene, *NCED3*, were higher in the OE lines after 1 h or 2 h of dehydration stress compared with the WT. Finally, the expression of *ABF3*, a key TF involved in the ABA signaling pathway (Kang *et al.*, 2002), significantly increased in the OE lines after 2 h of dehydration stress compared with the WT.

Finally, to investigate whether cuticular wax biosynthetic genes were induced by dehydration stress in grapevine, we examined the transcript levels of *KCS6* (VIT_14s0006g02990) and *LACS4* (VIT_02s0025g01410), which are homologous to *A. thaliana KCS6* (AT1G68530) and *LACS3* (AT1G64400), and found both to be induced by dehydration stress, peaking at 24 h (Supplementary Fig. S3).

## Discussion

Early studies identified three bZIP-type ABRE-binding proteins, *AREB1*, *AREB2* and *AREB3* from *A. thaliana*, using an ABRE motif (Uno *et al.*, 2000), and *ABF3* and *ABF4/AREB2* were shown to play important roles in response to ABA and drought stress signaling (Kim *et al.*, 2004). *AREB1* enhances drought resistance in *A. thaliana*, involving the ABA signaling pathway (Fujita *et al.*, 2005). In addition, Furihata *et al.* (2006) showed that exogenous ABA activates the SnRK2 protein kinases, and that the activated SnRK2 proteins phosphorylate a Ser/Thr residue in the conserved domains (C1, C2, C3 and C4) of the downstream *AREB1* gene, allowing it to bind to the *cis*-acting ABRE element of downstream drought-related genes. The mechanism involving *AREB1* in the ABA core signaling pathway in *A. thaliana* is also present in economically important crops, such as rice. For example, ABA and drought stress can trigger a rice SnRK2 protein kinase to phosphorylate rice bZIP TF, *TRAB1*, a homolog of *A. thaliana AREB1* and *AREB2* (Kagaya *et al.*, 2002; Kobayashi *et al.*, 2005). Chae *et al.* (2007) found that another rice bZIP TF, *OREB1*, can also be phosphorylated by a SnRK2 family protein kinase, *OSRK1.* These studies suggest that similar regulatory responses to stress are evolutionarily conserved in plants.

In this study, we identified a group A bZIP TF, *VlbZIP30*, from grapevine. Phylogenetic analyses indicated that VlbZIP30 is most closely related to ABF/DPBF TFs of group A, with conserved domains predicted to contain phosphorylation sites (C1, C2, C3 and C4) (Fujita *et al.*, 2005), suggesting that the *VlbZIP30* may be involved in drought stress and ABA core signaling (Fig. 1; Fig. 2A). Further work is still required to verify whether the conserved domains (C1, C2, C3 and C4) of VlbZIP30 are phosphorylated and that the protein is involved in the ABA core signaling pathway.

Previous studies have demonstrated that overexpressing drought-induced genes in *A. thaliana* can cause hypersensitivity to ABA and increase tolerance to drought stress (Fujita *et al.*, 2005). In this study, overexpressing *VlbZIP30* in *A. thaliana* did not make the plants more sensitive to ABA, but increased their osmotic stress tolerance during germination and post-germination growth. Similar results were reported for *SAD1, AtTPS1, AtHD2C, CaXTH3, OsMYB3R-2* and *ABO3* (Xiong *et al.*, 2001; Avonce *et al.*, 2004; Cho *et al.*, 2006; Sridha and Wu, 2006; Dai *et al.*, 2007; Ren *et al.*, 2010). Given that *VlbZIP30* is a TF, its role in osmotic stress response is likely to involve regulating downstream gene expression. Indeed, a majority of the ABA- and drought-induced genes tested were induced in *VlbZIP30*-overexpressing *A. thaliana* following ABA and mannitol treatment. Ninety-five (38%) of the 248 genes identified (Fig. 5) have previously been identified and of these 54 were involved in abiotic stress, including stress marker genes (*RD20, RD22, RD26* and others), ABA core signaling components (6 PP2Cs: *ABI1, ABI2, HAI1, HAI2, HAB1, PP2CA*, and *PYL6)*, TFs (*ABF2, ABF3, ABF4* and others) (Table 1). Nine *A. thaliana* PP2Cs belonging to cluster A have been identified (Schweighofer *et al.*, 2004), and studies have shown that 6 of them (*ABI1, ABI2, HAI1, HAI2, HAB1*, and *PP2CA*) function as negative regulators of ABA signaling, with their mutants showing hypersensitivity to ABA during seed germination and seedling growth (Merlot *et al.*, 2001; Saez *et al*, 2006; Yoshida *et al.*, 2006; Bhaskara *et al.*, 2012). In addition, the *ABI1, ABI2, HAI1* and *HAI2* genes are known to act in a negative feedback regulatory loop of the ABA signaling pathway (Merlot *et al*, 2001; Bhaskara *et al.*, 2012). A sextuple mutant impaired in six PYR/PYL receptors was shown to be very insensitive to ABA during seed germination and seedling growth (Gonzalez-Guzman *et al.*, 2012). Here, we found that the transcript levels of the 6 PP2Cs were all significantly higher in the *VlbZIP30-overexpressing* lines in response to osmotic stress, while the expression of *PYL6* was lower (Table 1). Consistent with this, the OE lines were found to be insensitive to ABA. These results suggested that *VlbZIP30* may play a role in the ABA core signaling pathway under osmotic stress conditions, and be involved in a negative feedback regulatory loop of the ABA signaling pathway in *A. thaliana* during the seedling stage.

Previous studies have shown that guard-cell movement mediated by ABA is a primary mechanism to prevent water loss under dehydration stress conditions (Kang *et al.*, 2002; Fujita *et al*, 2005). In this study, we found the stomatal closure regulated by ABA to be impaired in the OE lines (Supplementary Fig. S5), but that the expression the stress-marker genes (*RD20, RD26* and *ERD10*) was not be significantly different between the WT and OE lines. However, the expression levels of cuticular wax biosynthesis genes (*MYB96, KCS6* and *LACS3*) were significantly up-regulated in the OE lines compared with WT plants at the adult stage under dehydration stress (Fig. 11E). In addition, the expression of *HB7* and two PP2C genes (*HAI1* and *HAI2)*, which are mediators of a negative feedback regulatory loop of the ABA core signaling pathway in *A. thaliana* (Bhaskara *et al.*, 2012; Valdes *et al.*, 2012), were up-regulated in the OE lines compared with WT plants under dehydration stress (Fig. 11E). Our data suggest that overexpression of *VlbZIP30* enhances the tolerance of *A. thaliana* to dehydration stress at the adult stage through regulating cuticular wax biosynthesis related genes in the ABA signaling pathway. This is consistent with previous studies (Seo *et al.*, 2011; Cui *et al.*, 2016; Lee *et al.*, 2016a).

The ABRE (PyACGTGGC) and G-box (CACGTG) elements were identified as bZIP TF *cis*-binding elements regulating gene expression in response to ABA and drought stress in many plants, including *A. thaliana* (Uno *et al.*, 2000), rice (Liu *et al.*, 2014) and wheat (Wang *et al.*, 2016). In this study, we identified 39 *A. thaliana* genes and 35 predicted grapevine genes (Supplementary Data S2) that may be directly or indirectly regulated by *VlbZIP30.* Seventeen (43.6%) of the 39 *A. thaliana* genes have been found to be involved in drought stress, including *RD26, AFP1/3, PP2CA, HAI1/2, ABF3, NAP, MYB74, WRKY28* and *PUB19* (Table 1), implying that our analytical methods and results are very credible. Other genes found here that have not been previously characterized may therefore also be involved in drought stress signaling. In contrast, there has been little characterization of the 35 grapevine genes, and only 3 were identified as being involved in ABA or drought stress signaling. *ABF2/bZIP39* (VIT_18s0001g10450), which has been associated with both stimuli (Nicolas *et al.*, 2014; Tu *et al.*, 2016a), can transiently transactivate the expression of *NAC17* (VIT_19s0014g03290) and *PUB19* (VIT_17s0000g08080) in response to ABA treatment (Nicolas *et al.*, 2014; Pilati *et al*, 2017).

A perfect *A. thaliana* G-box (CACGTG) and a putative grapevine G-box (MCACGTGK) element were significantly enriched in the promoter of the 39 *A. thaliana* genes and 35 predicted grapevine genes (Fig. 9B, D). The highly conserved G-box motif (CACGTG) is regulated by bZIP TFs in organisms ranging from yeast to humans (Ezer *et al*, 2017). Ezer *et al.* (2017) constructed an available gene expression network (www.araboxcis.org) for prediction of genes regulating the G-box, or a set of genes regulated by the G-box. They identified approximately 2,000 seedling-expressed genes expressed in 229 RNA-seq samples of 7-d-old *A. thaliana* seedlings that are highly likely to be regulated by a perfect G-box motif (CACGTG) in their promoter, and predicted how bZIP proteins might regulate these genes. These results suggest that *VlbZIP30* is likely to enhance *A. thaliana* drought tolerance by regulating downstream genes containing a perfect G-box (CACGTG). Large-scale transcriptome analyses also show that the G-box (CACGTG) was highly enriched in stress-responsive genes in grapevines (Wong et *al.*, 2017). These results suggest a general conservation in promoter framework, gene expression dynamics and gene regulatory networks. We also used two different grapevine-related databases to gain support for the potential roles of the 35 grapevine genes in ABA and drought stress.

We noted that 74% and 84% (a total of 27) candidate genes were significantly up-regulated under ABA or drought treatment, respectively (Fig. 10A, B), and that the expression of some of these genes was up to 64-fold induced (Fig. 10B). In addition, we found by qRT-PCR analysis that the expression levels of 16 randomly selected genes from the 27 genes (including *VvPP2C9*, *VvPP2C37* and *VvABF2*) were significantly up-regulated by ABA or dehydration treatment (Fig. 10C; Supplementary Fig. S3), suggesting that the 27 candidate genes may involve in ABA or dehydration stress in grapevine. These results suggest that *VlbZIP30* may be involved in drought stress signaling in grapevine via regulation of the 27 grapevine genes containing the grapevine G-box (MCACGTGK). This conclusion is supported by the observation that 17 of the 39 *A. thaliana* homologous genes have previously been found to be involved in drought stress. Further studies be required to elucidate the functions of these regulated ABA and drought stress regulated grapevine genes.

## Supplementary data

Fig. S1. *VlbZIP30* mRNA levels in wild-type (WT) and transgenic plants analyzed by qRT-PCR.

Fig. S2. Phenotypes of wild type (WT) and *VlbZIP30* overexpressing transgenic lines at the seed germination stage under mannitol and abscisic acid (ABA) treatments.

Fig. S3. Gene expression profiles of selected *VlbZIP30-induced* grapevine candidate genes analyzed using qRT-PCR.

Fig. S4. Reactive oxygen species (ROS) levels and oxidative enzyme activities in wild type (WT) and transgenic lines (OE).

Fig. S5. Stomatal closure in response to 10 μM exogenous ABA in 3-week-old wild type (WT) and transgenic lines (OE).

Table S1. Specific primers used for qRT-PCR. F, forward; R, reverse.

Table S2. Differentially expressed genes in the *VlbZIP30* transgenic plants (OE / WT) based on an expression level differences (FDR<0.05) of at least two-fold under control conditions from the transcriptome data.

Table S3. Selected genes involved in other biological processes based on expression level differences (FDR<0.05) of at least two-fold in the *VlbZIP30* transgenic plants under ABA or mannitol stress treatment from the transcriptome data.

Data S1. Grapevine homologs of the up-regulated genes identified in *Arabidopsis thaliana* OE lines compared with WT plants.

Data S2. Names and annotations of 35 grapevine genes and 39 *Arabidopsis thaliana* genes.

Methods S1. Vector construction.

## Author contribution

X. Wang and M. Tu designed the study. M. Tu and X.H. Wang contributed to the experiments, X.H. Wang and D. Wang constructed the vectors, Y. Zhu performed the qRT-PCR analysis, M. Tu and D. Wang performed data analysis. M. Tu, X.H. Wang, X. Zhang and Y. Cui performed transcriptome data analysis, Z. Li, Y. Li and M. Gao assisted with the data analysis. M. Tu and X. Wang wrote the manuscript. All of the authors approved the final manuscript.

## Acknowledgments

This work was supported by the National Natural Science Foundation of China (31572110), as well as the Program for Innovative Research Team of Grape Germplasm Resources and Breeding (2013KCT-25). We thank PlantScribe (www. plantscribe.com) for careful editing of this manuscript.

